# Split-trial analysis reveals the information capacity of neural population codes

**DOI:** 10.1101/2025.11.27.690979

**Authors:** Dylan Le, Xue-Xin Wei

## Abstract

Understanding how correlated neural noise affects neural population coding is a basic question in computational and systems neuroscience [1, 2, 3, 4]. Recent theoretical work suggests that shared noise along the stimulus encoding direction is the primary factor that limits information encoding (i.e., information-limiting noise) [5, 6]. Despite this theoretical insight, it has been difficult to test it experimentally due to the challenges in inferring information-limiting noise from neural data. To overcome this challenge, we have developed a method (*i.e*., split-trial analysis) to partition the noise in a neural population into information-limiting noise vs. non-information-limiting components. Our method is simple to implement, yet it is highly effective given a limited amount of data. Results from extensive numerical simulations show that split-trial analysis substantially outperforms existing methods in accuracy, efficiency, and robustness. Applications of split-trial analysis to a number of neurophysiological datasets reveals insights into the precision of the neural codes for several systems. First, it reveals a substantial amount of information-limiting noise in the mouse head direction system. Second, it uncovers a small yet positive information-limiting noise in the orientation code in mouse V1. Third, we discover that the information-limiting noise in the macaque pre-frontal cortex is highly consistent over time during a simple saccade task. Split-trial analysis is a general technique that should be widely applicable in analyzing the properties of neural population codes.

## Introduction

Neural noise is correlated among neurons [1, 2, 7]. Noise correlations (NCs) can vary depending on task conditions [3, 8, 9, 10, 11], and may inform network computations [4, 12, 13, 14]. One important question has been whether the presence of NCs would benefit or diminish neural coding capacity [15, 2, 16]. Somewhat confusingly, earlier work suggests that NCs can increase or decrease coding capacity depending on the detailed structure [2, 1, 15, 17, 18, 19, 20, 21]. A unifying theoretical view begins to emerge from recent work [5, 12], that is, the key limiting factor for coding capacity is the shared noise along the stimulus encoding direction (which we will refer to as the *information-limiting noise*).

Despite this important theoretical insight, it remains difficult to measure information-limiting noise experimentally, due to its potentially small magnitude and the presence of other forms of non-information-limiting NCs [5]. While some recent studies reported substantial information-limiting noise in neural systems [22, 23, 24, 25, 26], others have found the opposite [27]. One existing approach relies on empirical estimates of Fisher information, then extrapolating to infer the information limit of large populations [23, 25]. However, direct estimation of Fisher information [28] requires estimation of the noise covariance matrix, which is difficult given the limited number of trials in most experiments. An alternative approach is to decode the stimulus and then use the Cramer-Rao bound [29] to convert the magnitude of the decoding error into an estimate of Fisher information [5]. However, prior methods require a large number of neurons and trials, and thus are severely limited in practice.

To address this problem, we have developed a method for inferring information-limiting noise based on splitting an individual trial into multiple sub-trials. This split-trial analysis method creates multiple repeated measurements within a single trial so that one can dissect (i) the noise shared by all neurons (a proxy of information-limiting noise) and (ii) the noise private to a subset of neurons. Numerical simulations reveal that split-trial analysis substantially outperforms existing methods in its sample efficiency, accuracy, and robustness. Applications of split-trial analysis to neural population recordings reveal a substantial amount of information-limiting noise in the mice head direction system, and a small amount of information-limiting noise in mice V1. It also reveals the temporal structure of information-limiting noise in the macaque prefrontal cortex during a simple saccade task.

### Accurate and efficient inference of information-limiting noise using split-trial analysis

The key idea of the proposed *split-trial analysis* is to create repeated measurements within a single trial by splitting the neural population into non-overlapping subsets. By decoding the neural activity from the two splits, one can partition the errors into those caused by shared noise across the two splits and the noise private to each split (Fig. 1a). Concretely, suppose that a stimulus variable *θ* is encoded in a large population of neurons from which we were able to measure the response of 2*N* neurons. We first split the neural population into two halves, then perform decoding analysis on each split (*e*.*g*., using Bayesian decoding). For a given stimulus *θ*, denote the decoded stimulus from the two splits as 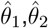, respectively. Intuitively, if the decoders across the two subsets are perfectly correlated, the decoder constructed based on all neurons will behave identically to the decoder based on each subset, thus doubling neurons used in decoding would not change the estimate or reduce the decoding errors. This implies that the decoding errors come from information-limiting noise. In the other extreme, if the estimates from the different splits are independent, there is likely no information-limiting noise– if there were, such shared noise would lead to positive correlation in the decoding errors of the two splits. Thus, we expect that the corelation of the decoders from the splits should reflect the magnitude of the information-limiting noise. Formally, through mathematical analysis (see Methods and SI Sections B & C), we find that the information-limiting noise can be estimated from a surprisingly simple estimator, *i*.*e*., the covariance of the two decoders Cov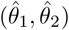.

**Figure 1:**
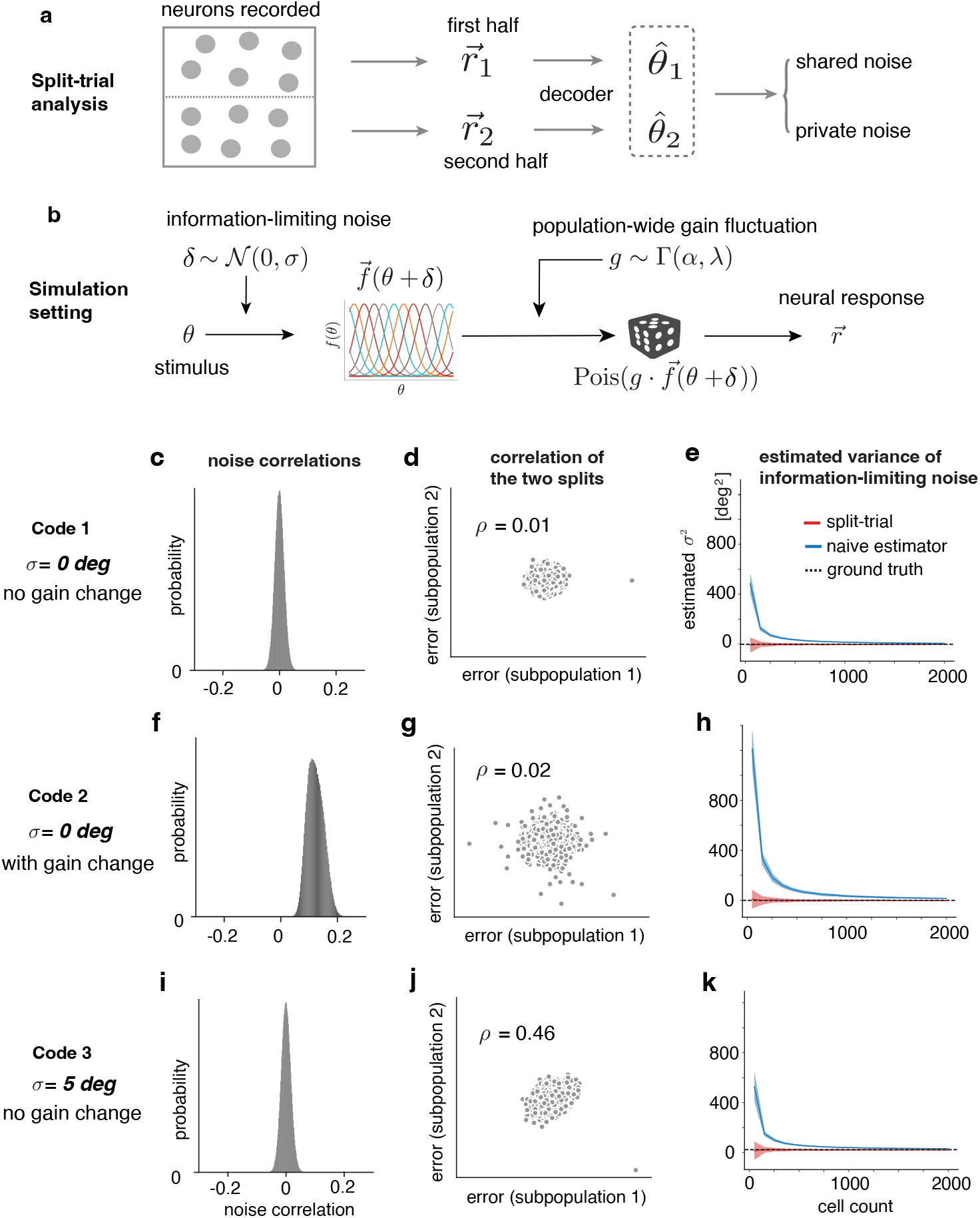
Split-trial analysis recovers information-limiting noise. **(a)** Overview of split-trial analysis. The neural population is first partitioned into two halves, which are both used to train individual decoders on the same set of trials. The decoding errors from both decoders are then leveraged against each other to separate shared (information-limiting) noise from noise that is private to each split. **(b)** Simulation pipeline for evaluating the methods. **(c**,**d**,**e)** In this case, we construct a neural population code (Code 1) based on tuning curves and Poisson spiking noise. **(c)** shows the distribution of the noise correlations, which is centered around 0. **(d)** shows the scatter plot of the two decoders from the two splits. **(e)** The inferred variance of the information-limiting noise for split-trial analysis and a naive estimator based on direct decoding. The ground truth information-limiting noise is 0 in this case. **(f**,**g**,**h)** Similar convention as **(c**,**d**,**e)**, but for a neural population code (Code 2) specified by tuning curves, Poisson spiking noise, and a population-wide gain fluctuation. No information-limiting noise is added. **(i**,**j**,**k)** Similar convention as **(c**,**d**,**e)**, but for a neural population code (Code 3) specified by tuning curves, Poisson spiking noise, and non-zero information-limiting noise (*σ* = 5 deg) added through corrupting the stimulus.

We illustrate split-trial analysis by simulating and analyzing three neural population codes (Fig. 1b). Code 1 is a standard neural population code with independent Poisson spiking noise. For Code 2, population-wide gain fluctuations [30, 31] (which is non-information-limiting) was added, that is, the response gain of neurons increases or decreases together on a trial-by-trial basis. For Code 3, information-limiting noise was added by injecting noise (*σ* = 5 deg) to the stimulus. Despite that both Code 1 and Code 2 have no information-limiting noise, their NCs are markedly different (comparing Fig. 1c and Fig. 1f). In contrast, Code 1 and Code 3 have different information-limiting noise, yet their distributions of NCs are similar (comparing Fig. 1c and Fig. 1i). These results show that the magnitude of NCs may not determine the information-capacity of a neural population code, consistent with the results in [5]. Crucially and central to our claim, these simulation results also suggest that the correlation of the decoders from the two splits strongly reflects the magnitude of the information-limiting noise (Fig. 1d,g,j). When information-limiting noise is absent (Code 1 and Code 2), the two decoders are largely uncorrelated. In the presence of information-limiting noise, the two decoders exhibit a strong positive correlation (Fig. 1j; also see Fig. S1 and Fig. S2).

Using simulated responses from these codes, we next benchmark the performance of our split-trial estimator against an existing method proposed in [5], for which the information-limiting noise is estimated based on a cross-validated decoder. We find that, for cases with zero information-limiting noise, estimates from the split-trial analysis are generally centered around 0 (Fig. 1e,h). Estimates from split-trial analysis can be smaller than 0 – this is a feature of our method and is important for an unbiased estimate of the variance of the information-limiting noise. In contrast, the estimated noise based on existing method is consistently larger than 0, and is thus heavily biased. When the information-limiting noise is non-zero (Fig. 1k and Fig. S2), our method is able to accurately recover its magnitude.

While the variance of information-limiting noise is of interest, for some applications, it may also be useful to infer its entire distribution, based on which various statistics (such as median absolute error, standard deviation) can be extracted. This may be particularly relevant when the distribution of information-limiting noise is heavy-tailed. We further developed a method to extract the entire distribution of information-limiting noise based on a deconvolution approach (see Methods and Fig. S3).

#### Generalization to binary estimation tasks

In addition to continuous estimation tasks, information-limiting noise have been studied in binary classification tasks [22, 23], a classical paradigm in psychophysics and neuroscience [32]. Suppose that we observe the neural activity corresponding to a pair of stimuli *θ*_1_ and *θ*_2_. One can ask how well one can discriminate the two stimuli (i.e., d-prime) from the response of a large neural population. The split-trial analysis can be generalized to this setting. The idea is to split the population into two sub-populations, and then apply a classifier to each sub-population to extract the decision variables for classification. These decision variables play similar roles as the decoders in the continuous case described earlier. We develop techniques that use the covariance of the decision variables from the two splits to estimate the information-limiting noise.

We benchmark the performance of our method and three existing methods [5, 23, 22] for estimating the magnitude of information-limiting noise in binary-classification tasks. The first baseline method [5] provides a naive estimate of the d-prime using linear classifiers with all neurons available. The second method [22] first performs dimensionality reduction, and then calculates the noise covariance, based on which the d-prime is estimated. The third method [23] directly estimates the generalized Fisher information based on noise covariance, and then relies on extrapolation to infer the information-limiting noise. Based on extensive numerical simulations by varying the number of trials, neurons, and the ground truth d-prime values, we find that the split-trial analysis can more efficiently and reliably recover the ground truth. Fig. 2 shows example results from these numerical simulations (more results based on a range of other sample sizes are shown in Fig. S4). When neural variability is dominated by large information-limiting noise (Fig. 2a,d), estimating information-limiting noise is relatively easy. All four methods (including the naive method) perform well when d-prime is equal to 2. As the information-limiting noise becomes smaller, the problem becomes more difficult. For these more challenging cases, only our method can reliably infer the information-limiting noise with a sample of 500 per stimulus. The naive method [5] and dimensionality-reduction method proposed in [22] substantially underestimate the magnitude of information-limiting noise (Fig. 2e,f). The method based on extrapolation by [23] exhibits substantial instability depending on the number of trials and the exact values of ground-truth d-prime values (Fig. 2e,f; also see additional results in Figs. S5-S7). For much smaller sample sizes, split-trial analysis also under-estimates the d-prime values, but to a lesser extent compared to other methods.

**Figure 2:**
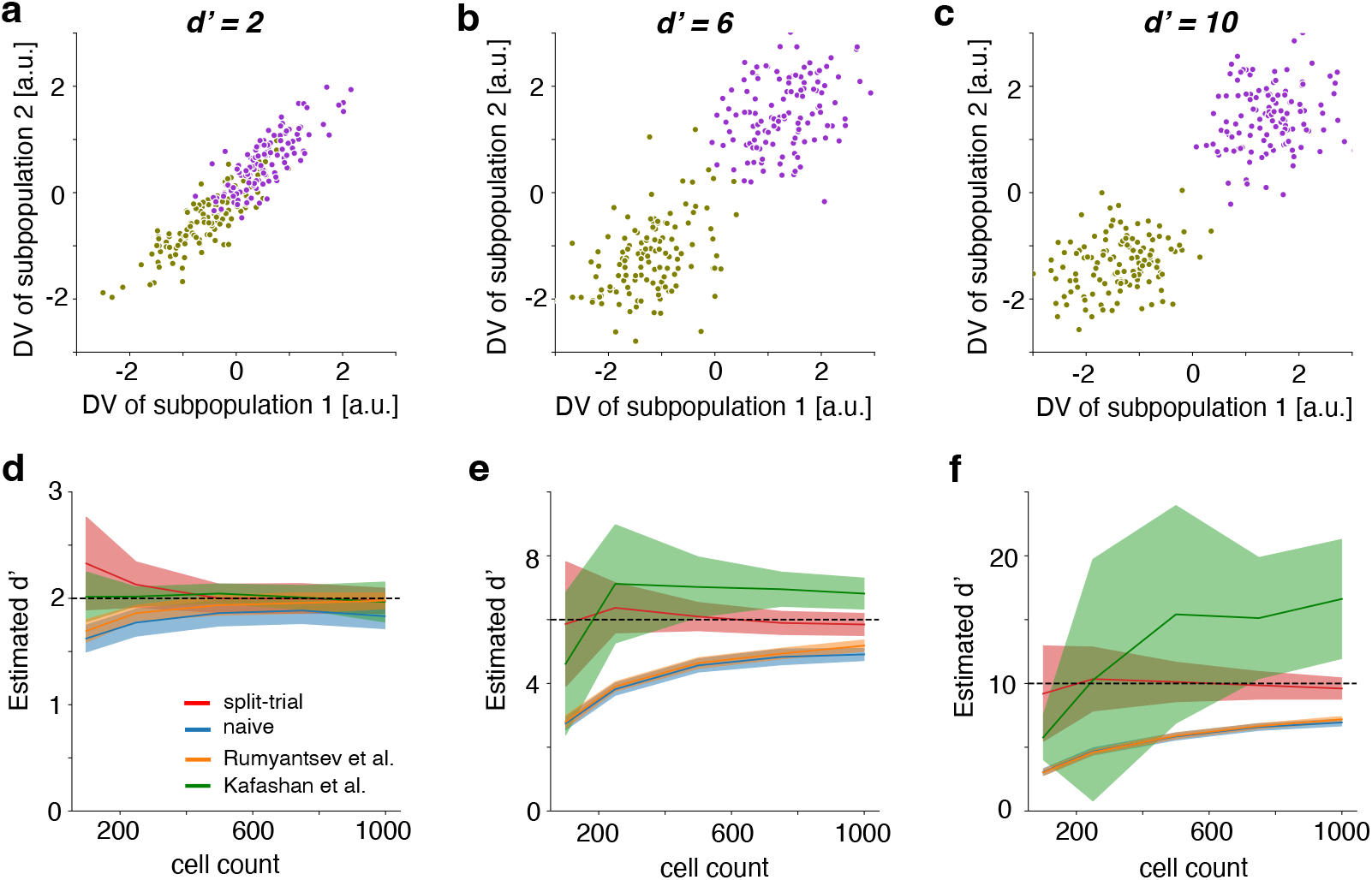
Benchmarking the performance of split-trial analysis in recovering the ground truth d-prime in binary classification tasks. **(a)** Scatter plot of the decision variable (DVs) for classifiers trained on two separate subpopulations of simulated neural datasets when d-prime =2. The distributions for each class are shown in gold and purple, and it can be observed that in higher noise (low d’) regimes, correlation between the two splits is more strongly observed. **(b**,**c)** Similar to **(a)**, but for d-prime =6, 10, respectively. **(d)** We estimated d’ in simulated data using a variety of different approaches to estimate information limiting noise, using 500 samples/stimulus, when d-prime =2. **(e**,**f)** Similar to **(d)**, but for d-prime = 6 and 10, respectively.

### Application 1: information-limiting noise in head direction system in mice

We first apply our method to study the head direction system [35], which encodes the brain’s internal compass in multiple species [36, 37]. The head direction system has been modeled using continuous attractor networks [38, 39, 40]. This type of model predicts that neural activity forms an activity bump and that neural noise manifests itself as coherent shifts along the ring manifold (Fig. 3a), thus predicting the existence of information-limiting noise [5] in head direction system. To test this prediction, we re-analyzed a recently published calcium imaging dataset that simultaneously recorded 100-200 head direction cells simultaneously in free-moving mice [34]. Prior results based on these data showed that head direction can be decoded with good precision [34]. However, it remains unclear whether the remaining decoding errors is due to information-limiting noise or non-information-limiting noise from limited numbers of neurons sampled. This question is challenging to address given existing techniques and sample sizes. Our method provides a unique opportunity to address this question.

**Figure 3:**
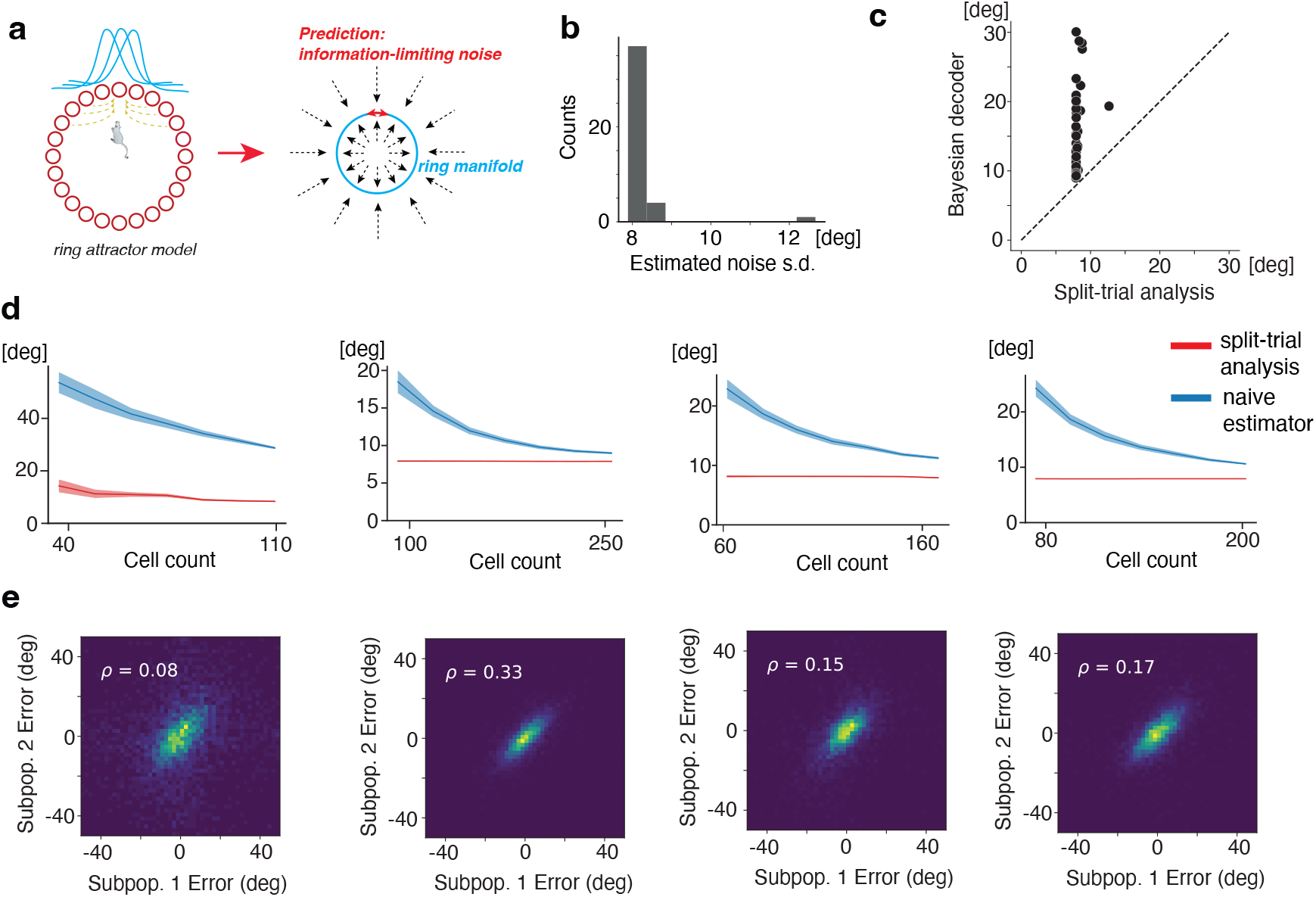
Results on head direction cells. Data reanalyzed from [33].**(a)** According to the ring attractor model commonly used to model head-direction cells, noise in the internal compass of the animal should manifest in a way identical to information limiting noise. **(b)** Histogram displaying the estimated information-limiting noise *σ* over 42 head direction population recording sessions from [34]. Decoding was done using a ZIG decoder as in the original work, and we estimate a mean information-limiting *σ* of ≈ 8.1 deg. **(c)** Plotting the estimated information limiting *σ* on the x-axis, and the estimate from a naive decoder (such as a Bayesian one) on the y-axis, we observe that our technique not only reports a generally lower information limit, but also that the variance of our estimator is substantially lower for this system. **(d)** Scaling in estimates of information-limiting *σ* as cell population size varies in four example sessions for both the split-trial analysis and a naive estimator. **(e)** 2D histograms of the error density across two subpopulations for a single iteration used to generate the plots in **(d)**, with Pearson’s correlation value noted.

Using a recently developed Bayesian decoding method for calcium imaging data [41], we first quantify how decoding errors vary with the number of neurons used in the decoding analysis by sub-sampling neurons. For most sessions, increasing the number of neurons used leads to a smaller error (Fig. 3d). Even when the number of neurons approaches the largest population size recorded in individual sessions, there does not appear to be a clear sign of saturation in the mean squared decoding error. These results suggest that inferring the magnitude of information-limiting noise using standard decoding analysis is difficult.

We then use the proposed split-trial analysis to analyze these data. We split the recorded neural population from each session into two halves, and perform decoding analysis on each of the two subpopulations. We find that the decoding errors between the two splits are only modestly correlated (Fig. 3e), which implies that the decoding error would be further reduced with a larger population size. Using the covariance of the decoding errors from the two splits to estimate the information-limiting noise, we find that the standard deviation of the estimated information-limiting noise is ∼ 8.15 deg (s.d. = 0.74 deg across sessions). Fig. 3b shows a scatter plot of the decoding errors inferred from the standard decoding approach and the information-limiting noise estimated from the split-trial analysis. The results show that, while there is substantial variability in the decoding error across different sessions, the inferred information-limiting noise is highly consistent across sessions. Furthermore, within individual sessions, we find that this value is also highly consistent when the number of neurons used is larger than 50. This observation further illustrates the robustness of our results. In addition, we used the deconvolution approach to directly estimate the full distribution of the information-limiting noise. We find that the estimated MAE is 4.4 deg, which is smaller than the MAE from a Bayesian decoding analysis (∼ 6 deg; [33]) The magnitudes of inferred MAE from split-trial analysis are also highly consistent across sessions (s.d.=0.61 deg). Together, these results provide strong evidence for the existence of information-limiting noise in the head direction system. Such information-limiting noise is consistent with the predictions from continuous attractor models [38, 40, 5].

### Application 2: The precision of orientation code in mouse V1

We next study a classic neural code, i.e., the orientation code in the primary visual cortex [42]. Recent studies have led to conflicting results regarding the existence as well as the magnitude of information-limiting noise in the orientation code of mouse V1 [22, 23, 27]. To further investigate this question, we apply split-trial analysis to the large-scale calcium imaging data reported in [27], which typically recorded more than 10,000 neurons simultaneously in individual sessions.

In one of the experiments conducted in [27], stimulus orientations were continuously sampled from the whole circle. We first applied the standard decoding analysis to these data. Consistent with the report in [27], the mean squared error of the decoded orientation continuously decreases as the number of neurons used in decoding increases, and goes to about 2 deg when all neurons are used (typically 10k-20k neurons). However, it remains unclear whether the decoding error has saturated. We sought to address this question by applying the split-trial analysis to these data while varying the number of neurons used (Fig. 4c). We find that, using 10% of all neurons recorded in individual sessions, the inferred standard deviation of information-limiting noise is about 1 deg. The magnitude of the inferred information-limiting noise is largely stable when more neurons were included in the analysis. Using all neurons, the inferred information-limiting noise is only slightly reduced. The inferred value of the standard deviation of information-limiting noise is highly consistent with the 5 sessions (mean= 0.82 deg, sd =0.05 deg;Fig. 4d).

**Figure 4:**
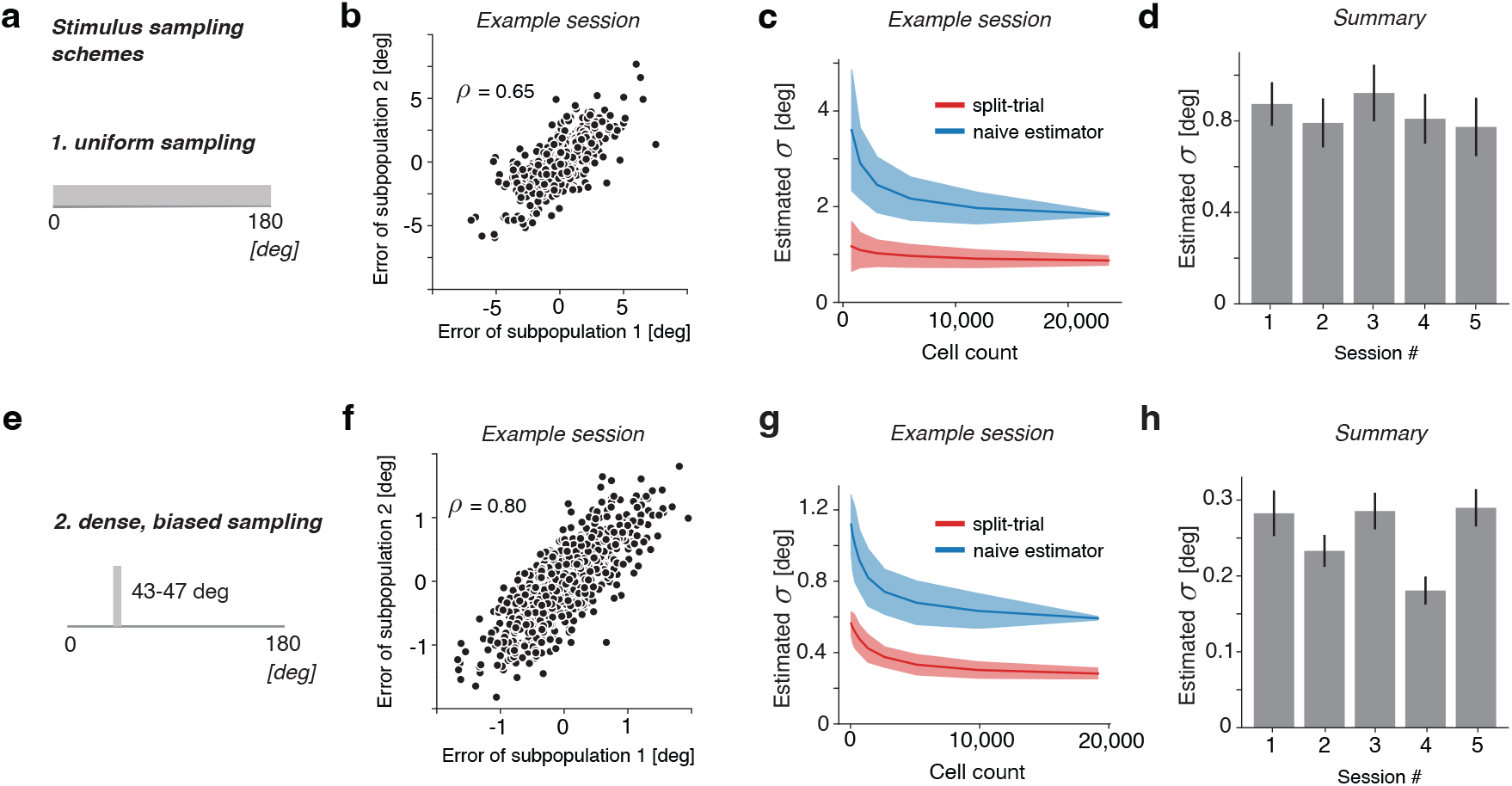
Results on the orientation code in mice V1 from two different experimental paradigms. Data reanalyzed from [27]. **(a)** Schematic of the uniform stimulus sampling method. **(b)** Example scatter plots of the subpopulation errors for sessions from the static orientation and dense discrimination datasets, respectively. The modest degree of correlation in both indicates the presence of information-limiting correlations in the considered datasets. Data from from [27]. **(c)** An example session’s estimated information-limiting *σ* plotted against population size for static orientation and dense discrimination, respectively. **(d)** The estimated information-limiting *σ* for the static orientation and dense discrimination datasets, taken at the full population sizes for each session. One session was not visualized due to its large error bar. See Fig. S10 for results for all 6 individual sessions. **(e-g)**. Similar to **(a-d)**, but for another dataset that densely sample the stimuli from a small range, i.e., [43,47] deg.

We further analyzed a second dataset collected in [27] using a different stimulus sampling paradigm. In this experiment, orientations of the gratings were densely sampled within a small range of 4 deg. Applying the split-trial analysis, the results suggest that the information-limiting noise is around 0.26 deg (sd= 0.04 deg). This is smaller than the magnitude of the decoding error inferred from the direct decoding approach in [27], but remains larger than 0. We notice that reliably inferring information-limiting noise is more challenging in this dataset, as evident in the slightly decreasing trend of inferred information-limiting noise beyond 5,000 neurons (Fig. 4e). This is likely due to the tiny amount of information-limiting noise in this data.

The results based on these two experiments provide evidence for the existence of a small amount of information-limiting noise in V1. Our inferred magnitudes of the information-limiting noise in mice V1 are substantially lower than those reported in [22, 23], which are generally larger than 5 deg. One reason for such differences might be the smaller cell counts used in [22, 23] (on the order ∼ 1000 neurons or less) together with the inaccuracy in the estimation of d-prime given the limited number of neurons. When we apply standard Bayesian decoding using cell counts that are comparable to [23] and [22] via subsampling the data from Stringer et al [27], the estimated s.d. of information-limiting noise is generally around 4 deg (Fig. S10), which is closer to the values reported in [23] and [22]. Another contributing factor could be the different paradigms used in estimating information-limiting noise. In particular, the d-prime values computed based on discriminating a pair of well separated stimuli may be difficult to link to the s.d. of information-limiting noise from continuous estimation tasks when the neural response manifold is highly curved [43].

### Application 3: Temporal integration of stimulus information in frontal cortex

Neural computation involves the integration of information both over neurons and over time. As most previous studies examined correlated neural noise focused on the effect of integrating information over neurons in a fixed time window [15, 5], the degree to which temporal integration affects information encoding remains an important open question [44]. While noise correlations can be computed for neural activity across time, it remains unclear how they affect temporal integration for large neural populations. For simplicity, theoretical studies of neural population coding typically assume independent noise for different time windows, which implies a large benefit of integrating over a longer period. Empirical studies have examined the effect of temporal integration for individual neurons, and found that integration over several hundred milliseconds remains beneficial [45, 46]. However, results obtained from individual neurons or small neural populations may not generalize to large neural populations. For a large neural population, the impact of temporal integration on the encoded information should mainly depend on the temporal dynamics of information-limiting noise. Consider the extreme case that information-limiting noise for two adjacent time windows is perfect correlated. In this case, integrating over these two time windows would lead to no improvement in information encoding. Conversely, if information-limiting noise at two adjacent timesteps is uncorrelated, temporal integration would substantially increase the information. So far, the temporal structure of information-limiting noise remains an open question.

We investigate this question in the context of monkey prefrontal cortex (PFC) when the monkeys were performing a simple left/right saccade task [25]. Briefly, on each trial (Fig. 5a), the monkey was tasked with fixating on a point in the center of a screen for a variable fixation period (400-800ms), after which the fixation point would disappear and a stimulus point would appear in either the animals’ left or right visual field. The animal would then be required to fixate for 500ms on the stimulus point, before a reward was stochastically delivered on correct trials. Up to 800 neurons in PFC were recorded simultaneously in individual animals during the experiments (n=4 sessions from two monkeys). We focus on neural activity during the 300ms before the reward period. Performing standard decoding analysis on all neurons recorded, we find that the d-prime values are close to 7 (mean =6.82, s.d.= 0.8), consistent with the original report in [25]. Examining the correlation of the decision variables across the two splits, we find that they are only modestly correlated, which implies that even larger populations may lead to even larger d-prime values. Applying the split-trial analysis to the neural population response in the 300ms window defined above, we find that the inferred d-prime values for the four experimental sessions are 13.3, 13.5, 10.2, and 11.2, respectively. These values are substantially larger than those from the standard binary classification analysis. Meanwhile, they are substantially smaller than the values inferred in [25] using a method based on extrapolation. The inferred d-prime values are consistent for the two sessions of individual monkeys (Fig. 5c), with monkey V exhibiting larger d-prime values compared to monkey W.

**Figure 5:**
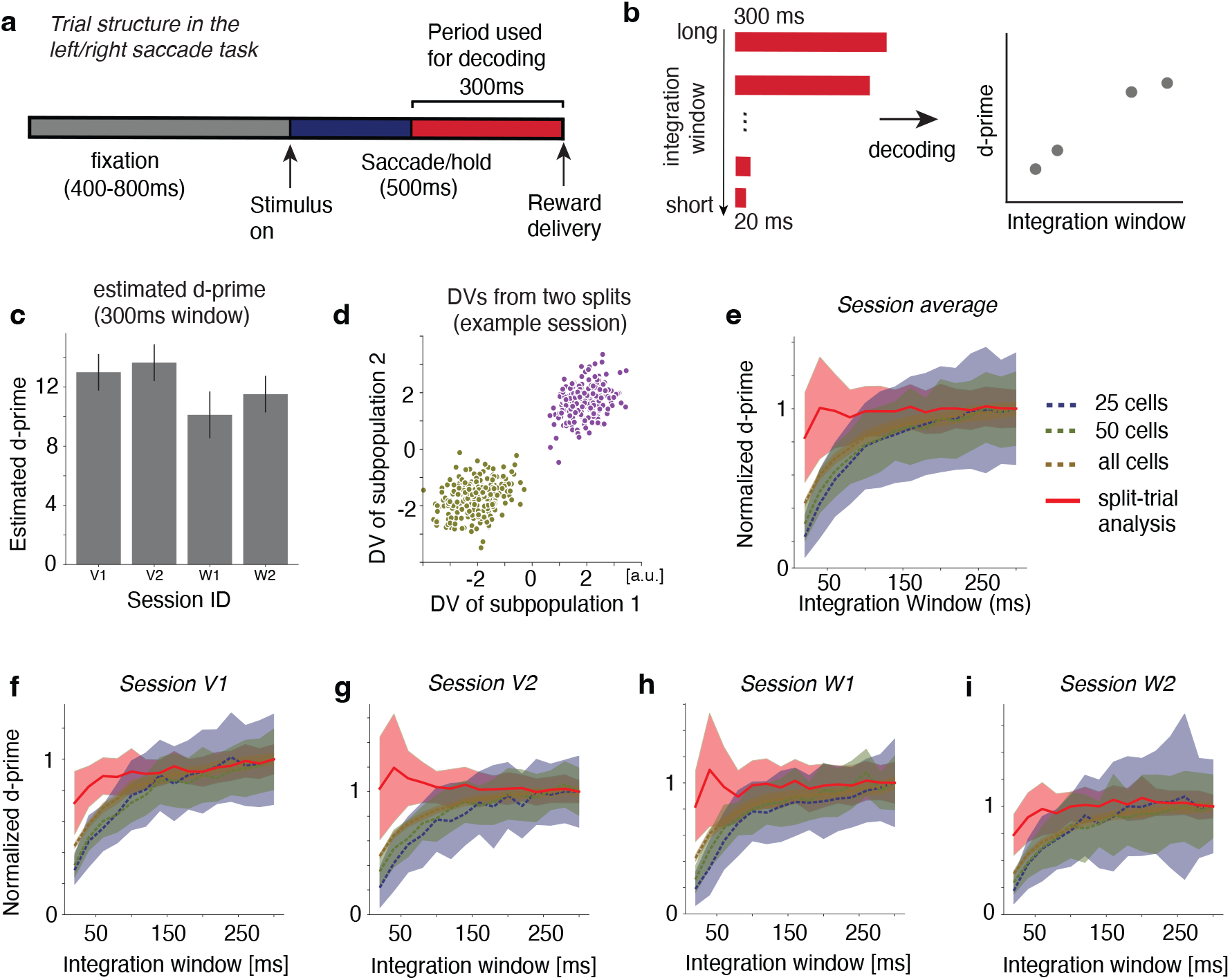
Results on the neural code in macaque PFC during a left/right saccade task. Data reanalyzed from [25]. **(a)** Schematic of the structure of an individual trial. **(b)** Schematic of the analysis procedure. We gradually decrease the length of the time window used for the analysis. For each time window, we inferred the d-prime value based on split trial analysis and the standard classification analysis (naive approach). **(c)** Estimated d’ limit for PFC recording sessions from Bartolo et al., dataset consisted of two animals (labeled ‘V’ and ‘W’) across recording sessions for each animal. **(d)** Example DV scatter for a trial from session V1. **(e)** d-prime values in the limit of infinite number of neurons was estimated using both the split trial analysis (solid lines) and the naive approach (dashed lines) for increasingly large spike integration windows for the time period starting 300ms before reward delivery on successful trials. Each plot shows change in d’ limit as integration window is widened across different total population sizes (colors in legends), normalized by the mean value at the widest integration window. The curves in this panel indicate that, in this dataset, longer integration times do not substantively change the split trial method’s estimation of the information limit of the system in this task, as is the case for the naive approach. This indicates a strong and stable signal of information can be recovered in PFC with as little as 20ms spike data from the split analysis approach. **(f-i)** Individual plots for each session, in the same manner as panel **(e)**.As was the case in panel **(e)**, curves displayed estimated d’ using 25, 50, and all cells recorded in the session.

We next turn to our key question of how correlated noise affects information during temporal integration. We first applied the standard decoding analysis on neural populations of different sizes, while varying the time-window of the decoding. We tested a number of time windows, ranging from 300ms to 20ms with a 20ms decrement (Fig. 5b). We find that for small neural populations (e.g., n=25 cells), d-prime increases substantially with increasing time window. When n=25, the d-prime values obtained at 300ms are on average 2.3 times larger compared to those from 20ms. As the number of neurons increases, the benefit of temporal integration reduces. Applying the split-trial analysis to all neurons recorded in individual sessions, we find that there is a benefit of temporal integration below 40ms. However, beyond 40ms, there is little improvement in the information. On average, the d-prime values based on a 40ms window are almost as large as those obtained from the 300ms window (Fig. 5e).

Because the analysis above involves a gradually changing time window, one possible explanation is that the saturation of information with larger time windows is due to the later period containing little or no information about the task variable. We perform additional analysis to rule out this possibility. Specifically, we use split-trial analysis to infer the d-prime values from each of the fifteen 20ms time windows during the 300ms period we analyzed. We find that there is still substantial amount of information in the later periods, although the information decreases modestly (Fig. S15). Together, these results suggest that, for this simple saccade task, information-limiting noise in PFC for the saccade location is highly correlated over time within individual trials. Given a sufficiently large neural population, each of the 20ms time window contains highly redundant information. One potential implication of this finding is that, for this simple saccade task, if the downstream areas have access to a large population of neurons, integrating the response from PFC over a brief time period is sufficient to transmit most information about the saccade location.

## Discussion

We developed a split-trial analysis method to infer the magnitude of information-limiting noise (or equivalently, the information capacity) for large neural populations. Split-trial analysis creates repeated measurements within a single trial, and it enables the partition of the noise of the neural population (captured by the decoding errors) into shared noise across the two splits and private noise specific to each split. Our results show that the estimated shared noise across two splits provides a surprisingly simple yet effective estimator of the information-limiting noise. While in this paper we have focused on information-limiting noise, the method can also be used to estimate the contribution of non-information limiting noise as a by-product for a finite neural population, which can be estimated by taking the difference between information-limiting noise and total noise.

The idea of splitting the population into subsets and performing decoding on each subset was mentioned in [27]. However, it was not recognized that it could be used as an effective way to infer information-limiting noise. Previously, it was proposed that the decoding errors of the two subpopulations should be generally correlated in the presence of correlated neural noise [27]. However, we have demonstrated that this is not the case. For example, while shared multiplicative gain fluctuations lead to large positive correlations in the neural population (Fig. 1f), the decoding errors from the two subpopulations are generally not correlated (Fig. 1g, Fig. S1). Importantly, we find that the correlation of the decoding errors across the two splits serves as a signature of the information-limiting noise, rather than arbitrary forms of NCs. This is a critical insight that was not appreciated previously.

The split-trial analysis does not suffer from severe estimation biases of previous methods. The standard method [5] based on directly decoding the stimulus overestimates the information limiting noise for a finite neural population. Our numerical results show that similar concerns apply to the method in [22], in particular when the information-limiting noise is small. Furthermore, our results show that a method based on extrapolation [23], while being well motivated, is unstable in practice. Split-trial analysis method is substantially more robust than these prior methods, and remains generally stable for different numbers of trials and neurons. One exception is that, when only a small number of trials are available to infer a large d-prime, split-trial analysis exhibits an under-estimation. Interestingly, we did not observe such biases for continuous estimation tasks. It is possible that a refined version of split-trial analysis may remove these biases when estimating d-prime in binary classification tasks.

Application of our methods to neurophysiological data reveals several new insights. First, the results revealed a substantial amount of information-limiting noise (s.d. ∼ 8 deg in standard deviation) in the mouse head direction system. The magnitude of this information-limiting noise is highly consistent in three mice we analyzed. This result is consistent with attractor dynamics, and provides an example of the existence of substantial information-limiting noise in spatial navigation system, in additional to recent work on place cells [26]. It remains to be tested whether head direction systems in other species exhibit similar amounts of information-limiting noise and whether the information-limiting noise depends on the experimental conditions.

The magnitude of the information-limiting noise inferred from mouse V1 data [27] is substantially smaller. When the stimuli were sampled continuously over orientation, the standard deviation of the information-limiting noise is slightly smaller than 1 deg. For the densely sampled dataset, the information-limiting noise is around 0.2 deg, yet remains larger than 0 deg. These values are substantially lower than those reported in two recent studies [22, 23]. Although the exact reasons of this discrepancy remain a subject of further investigation, we consider that the larger information-limiting noise inferred in [22, 23] could potentially due to a combination of the following: (i) a smaller number of neurons recorded and used in the analysis (10k v.s 1k); (ii) coarse sampling of the stimulus in their experimental design; (iii) an overestimation in the estimated information-limiting noise. Notably, we found evidence for a difference in the magnitude of the information-limiting noise when comparing the two experimental conditions in [27]. [27] considered that the difference in the decoding errors was due to the density of the trials sampled– an effect we also confirmed when downsampling the dense dataset. This difference may not be fully explained by the number of trials. When we subsample the second dataset (dense sampling) to match the number of samples in the first dataset (uniform sampling), we find that the magnitude of the inferred information-limiting noise is about 0.6 deg (Fig. S12), which remains generally smaller than the values inferred from the first experimental condition with uniform sampling of the stimuli. This raises the possibility that information-limiting noise may depend on the task conditions, such as stimulus statistics (i.e., sensory adaptation) and paradigms (i.e., binary classification v.s. continuous estimation).

Our method opens the door for studying the temporal dynamics of information-limiting noise. This was a difficult question to study due to the entanglement of information-limiting noise and spiking noise in neural responses. Applying split-trial analysis to study monkey PFC data collected during a saccade task shows that temporal integration in PFC has little effect beyond 40 ms when a large population is involved, unlike what was reported for the cases of individual neurons or small neural populations [46]. This suggests a large temporal correlation of the information-limiting noise, i.e., information-limiting noise is highly correlated over time. While our analysis approach is general, it is possible that this result may be specific to this simple task. For this task, being able to extract the information from a small time window would potentially be beneficial for making quick saccade planning. It would be interesting to apply our method to study whether temporal integration over longer timescales carries more substantial benefits in other decision tasks and other brain regions. In particular, for tasks that would benefit from longer integration and deliberation [47, 48], information-limiting noise may be less correlated over time.

One important yet under-investigated question is how the information-limiting noise depends on task conditions, e.g., attentional states [3] or learning [49]. Having multiple task conditions in a single experiment necessarily means fewer trials for each condition. The large number of trials required for inferring information-limiting noise under a given condition severely has limited the ability to task-dependence of information-limiting noise [50] in previous studies. With split-trial analysis, the ability to infer information-limiting noise reliably with much fewer trials should open up exciting opportunities for future experiments that investigate how network computations affect the capacity of the neural code and and temporal dynamics of information-limiting noise in neural circuits.

## Methods

### Simulations

For simulations, the tuning curve of an individual neuron for stimulus variable *θ* was assumed to be Gaussian (or von Mises function). For neuron *i*, the tuning curve is denoted as *f*_*i*_(*θ*), which is centered at the neuron’s preferred stimulus *θ*_*pref,i*_. The number of cells and tuning width were generally set for full coverage of the stimulus space. Neural spiking statistics were determined by the tasks that were being simulated (see below). Stimuli were sampled continuously over the stimulus range.

#### Continuous Estimation

When simulating neural activity responses for a continuous estimation task, the response was generated according to the following model

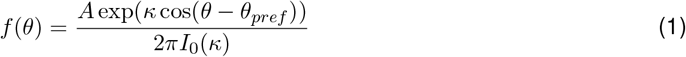

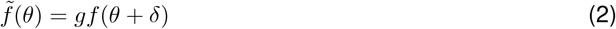

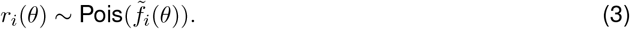

The neural tuning functions were modeled as von mises distributions centered at their preferred stimulus with parameter *κ* determining tuning curve width, parameter *A* controlling peak rate, and parameter *I*_0_(*κ*), the modified Bessel function of order 0, is essentially a normalizing factor that can be omitted for purposes of modeling neural tuning. The information-limiting noise was modeled as a shared input noise term *δ* ∼ 𝒩 (0, *σ*) which corrupted the input stimulus shared across the entire neural population (i.e. *δ* is the same for each neuron). For some simulations, a term *g* ∼ Γ(*α*, 1*/α*) is introduced to model the trial-by-trial gain fluctuation.

#### Discrete Stimuli & Binary Classification

When generating simulated data for binary classification tasks, the neural response was generated according to the following expressions:

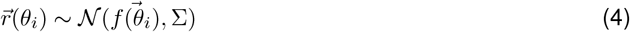

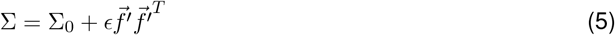

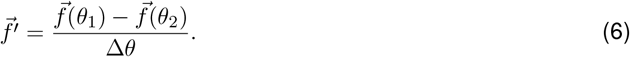

Matrix Σ_0_ is a covariance matrix for the population with no information-limiting component (i.e. a minimal projection along the 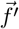 direction). This matrix was generated by constructing a random positive semi-definite matrix via Σ_0_ = *UDU*^*T*^, where *U* is a randomly drawn orthogonal matrix (in our case the *ortho_group* function from the scipy.stats package in Python), and *D* is a diagonal matrix determining the eigenvalues of Σ_0_. The entries of *D* were drawn from an exponential distribution with rate parameter one, but the largest values of D were multiplied by a factor of up to 5 to control for the level of noise correlations present in the neural population. Constructing Σ_0_ in this way enables one to manipulate non-information-limiting noise in the system directly by changing columns of *U* and entries in *D*. The parameter *ϵ* is the magnitude of the information limiting correlations present in the population.

### General data analysis pipeline

#### Continuous stimuli

For neural data (simulated or empirical) which came as a result presentation of continuous-valued stimuli, we used a naive Bayesian decoder to estimate the stimulus presented for each trial based on the neural population response [51], assuming neurons are noise independent. The decoded stimuli were compared to the ground truth to obtain the error for each trial. The decoding errors were used to perform the split-trial analysis.

#### Discrete stimuli

In the case of data generated from a discrete stimulus space, as in classical binary classification tasks, the neural response was used to train an linear support vector machine (SVM), although any well-trained classifier might suffice. Typically 70% of the data was held out for training and a 5-fold cross-validation procedure was used on the training data to obtain optimal model parameters. For the test data, rather than using the choice of decoded stimulus, we obtained the decision variables by taking the (1-dimensional) distance of each trial to the decision boundary from the trained SVM. These decision variables were then used to perform the split-trial analysis.

### Split-trial analysis

Split-trial analysis begins by partitioning our total population of *N* neurons into two subsets of *N/*2 distinct cells each. Then we train decoders on the data from each subset. Let 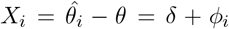 be the decoding error for both subpopulations. Here we provide a justification of the covariance estimator under a simple setting. For more rigorous mathematical justification, see Sections B & C of the SI. Assume that the decoders 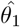 and 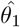 are unbiased, i.e., *E*[*ϕ*_1_] = *E*[*ϕ*_2_] = 0. Further assume that *δ* and *ϕ*_*i*_ are all independent. Intuitively, *δ* describes the shared (information limiting noise) term, and the *ϕ*_*i*_’s describe the private noise unique to the individual subpopulations. In fact, the random variables do not need to be gaussian as the following argument only requires they be mean zero, finite variance, and that they be independent, the assumption of Gaussian distribution was simply chosen for notational convenience. From this decomposition of the error we can calculate: *X*_1_ + *X*_2_ = *ϕ*_1_ + *ϕ*_2_ + 2*δ*; *X*_1_ − *X*_2_ = *ϕ*_1_ − *ϕ*_2_. Because *ϕ*_1_, *ϕ*_2_ and *δ* are independent, *Var*(*X*_1_ + *X*_2_) = *Var*(*ϕ*_1_) + *Var*(*ϕ*_2_) + 4*Var*(*δ*), and *Var*(*X*_1_ − *X*_2_) = *Var*(*ϕ*_1_) + *Var*(*ϕ*_2_). Thus, the variance of the shared noise (denoted as *σ*^2^) can be calculated as 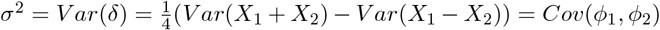. This result suggests that one can use *Cov*(*ϕ*_1_, *ϕ*_2_) to estimate the variance of information-limiting noise.

For a single iteration of the split-trial analysis, the neural population is randomly divided into two halves by assigning each neuron to one of two subpopulations. Then the decoding error will be considered *X*_1_, *X*_2_ for the first and second subpopulations, respectively. We then calculate *Cov*(*X*_1_, *X*_2_) as the estimator for the variance of the information-limiting noise obtained for a single iteration. Multiple iterations of split-trial analysis can be performed to get a distribution of the estimated variance of the information-limiting noise, from which the mean and the confidence interval of the estimated variance (or the standard deviation) of the information-limiting noise can be obtained.

### Discrete extension of split-trial analysis

We develop an algorithm to generalize split-trial analysis to the case of estimating d-prime values for binary classification tasks. Previous methods [22, 23] have also been developed to infer information-limiting noise based on binary classification. In this case, the decoding error 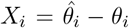 is no longer available. We find that the decision variables/logits (DVs) returned by the linear classifier for individual trials can be used similarly to decoded stimuli in the continuous estimation case. In this setting, the DVs are a measure of a given trials (signed) distance to the decision boundary, with sign determined by class label. According to the standard one-dimensional signal detection theory, one assumes that the decision variable for choosing the stimulus *i* follows a Gaussian distribution, *DV*_*i*_ ∼ 𝒩 (*µ*_*i*_, *ξ*), where *i* = 1, 2 for each subpopulation. Under this standard setting, d-prime can be defined as 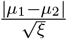. If these decision variables were directly extracted from a linear classifier, *ξ* would contain contribution from both information-limiting noise and non-information-limiting noise. The key idea is to use split-trial analysis to isolate the variability of the information-limiting noise.

Specifically, when extended to consider two subpopulations, we have twice as many distributions, but each DV can fall into only two distinct distributions. We then take the same decomposition as before to describe *DV*_*i*_ = *µ*_*i*_ + *ϕ*_*i*_ + *δ*, for a given subpopulation and trial, and a similar calculation as described previously enables us to leverage shared variability in both neural population’s DVs to estimate the shared information limiting noise in the decision space, *σ*. Let the empirical mean of the DVs for each split be *µ*_*i*_ = (*µ*_*i*1_, *µ*_*i*2_), where, for example *µ*_12_ would be the mean of the DVs corresponding to subpopulation 1, with trials where the stimulus presented was from class 2. In practice, *µ*_*i*_ is the empirical mean of the DV’s for each stimulus, e.g. 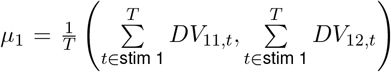. We first normalize both sets of DVs to have the same Δ*µ*_*i*_ = |*µ*_*i*1_ − *µ*_*i*2_| (typically normalizing them to both have *µ*_*i*_ = 1), reflecting our assumption that both splits are statistically identical. From there we can use the DVs as we treat ‘error’ in the continuous approach to obtain *σ*_*i*_. for subpopulation *i*, then 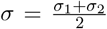. The information-limiting discriminability index (d’) is then calculated: 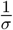, as we’ve previously normalized the distances between DVs representing different classes.

### Estimating the distribution of information limiting noise using a deconvolution approach

The distribution of the information-limiting noise may be non-Gaussian, in which case variance alone may be insufficient to describe the properties of information-limiting noise. We develop an approach that directly estimated the information-limiting error distribution. Let *X*_1_ *= δ +ϕ*_1_, *X*_2_ = *δ* +*ϕ*_2_ be the decoding error for the first split and the second split, respectively. Assume that *ϕ*_1_ and *ϕ*_2_ are independent, and furthermore their density functions are symmetric. It follows that *X*_1_ + *X*_2_ = 2 ∗ *δ* + *ϕ*_1_ + *ϕ*_2_, and *X*_1_ − *X*_2_ = *ϕ*_1_ − *ϕ*_2_. Because *ϕ*_1_ and *ϕ*_1_ are independent and symmetric, *ϕ*_1_ − *ϕ*_2_ and *ϕ*_1_ + *ϕ*_2_ follows the same distribution. Therefore, the probability density of *X*_1_ + *X*_2_ is the result of convolving the density of *X*_1_ − *X*_2_ and the density of 2*δ*. This suggests that density of 2*δ* can be estimated by solving a deconvolution problem [52]. Note that, for this deconvolution approach, it is important that the density of the decoding error is symmetric. This assumption of symmetric error distribution is not required when using the covariance of the decoders from the two splits to estimate the variance of information-limiting noise.

In our implementation, we assume that the density of 2*δ* has the form of 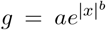, which includes Gaussian density function as an special case. The problem is to find a filter *g* for which *g* ∗ (*X*_1_ − *X*_2_) = *X*_1_ + *X*_2_. To find the best fit for the filter *g* we did a grid search over *a* and *b* and fit with least squares, using the empirical distribution *X*_1_ + *X*_2_ and *X*_1_ − *X*_2_ computed based on the decoding errors of the two splits. The range of exponents, *b*, and the range of widths, *a*, were continuously sampled in the range (0, 3], and (0, 5], respectively. After finding the best filter, we treated it as a PDF for the information-limiting noise distribution and extracted the standard deviation *σ* and the median absolute deviation to characterize the noise for the dataset.

### Dataset specific details

#### Ca^2+^ Imaging Dataset

In this study [27], calcium imaging was done in V1 for awake, head-fixed mice while they observed static oriented gratings whose orientations continuously spanned the circle. Analysis of the dataset followed the procedure laid out in the original author’s report. In short, we performed decoding using the neural activity collected while animals observed a collection of continuously distributed oriented static gratings. This decoding was done using their so-called ‘super-neuron’ based decoding procedure for both subpopulations. 48 super-neurons were spaced evenly across the range [0, 360] deg. Information limits were estimated using the deconvolution variant of the split analysis described below, as well as with the naive decoders trained from the original report from [27].

#### Head Direction Cell Dataset

In this dataset, calcium imaging was performed in the anterodorsal thalamic nucleus (ADN) of mice. The animals were kept in a chamber with a 360 deg LED screen. Vertical stripes were displayed at different regions of this screen during the baseline recording session, which is the period of the session that our analysis focused on. Our analysis pipeline for the head direction cell data from [34] also followed the original approach with small tweaks to fit our splitting procedure. The recorded responses from the head direction cells of the animal while it tracked the rotation of a moving landmark were fed into a ZIG decoder [41], as was done in the original paper by [34]. In fact, for each iteration, we trained two ZIG decoders, based on the activity of either of our two population splits. The head direction errors from both decoders for each trial were then used for split-trial analysis as previously described.

#### Macaque PFC Dataset

For the dataset from [25], animals were trained to fixate to a target on the center of a screen for as long as the fixation point stays on the screen (400-800ms). A stimulus would then appear on either the right or left of the animal’s visual field and the animal was expected to saccade to the point and hold its gaze for 500ms on the stimulus, after which a reward would be delivered 70% of the time. For our analysis, we trained an SVM on the neural activity in PFC from the first 300ms of the 500ms period preceding reward delivery or trial cessation. More specifically, we took the time a trial ended (due to either reward delivery or timing out) and considered the 500ms preceding that point to be the period when the decision was being encoded in the PFC. We then trained our SVMs on the first 300ms of that 500ms interval. The reason for this delineation is that it was often the case that animals could fixate on the stimulus for a period of time less than 500ms before saccading back to the stimulus and holding gaze for the required time. In parsing the data this way were were able to find a stable point of comparison across all the experimental trials. From the SVMs we recovered the decision variables as the linearized distance between the trials and the separating hyperplane. In this case, the ‘errors’ used for the split-trial analysis are the decision variables.

In order to assess the dynamics underlying information integration in PFC for this task, we also performed the split-trial analysis using data with increasingly wide time windows from the same starting time point. In this case we started with a 20ms wide window, estimating the information-limiting noise from the first 20ms of activity from our starting point. The window was increased by increments of 20ms up to the full 300ms wide window of activity used in the previously described analysis.

### Implementation of alternative methods

The PLS reduction technique used in [22] was reimplemented following the description from the original paper and from their own original code. In short, neural response data was split by trial into thirds, where one third of trials were used to train a PLS Regression model, one third (*r*_*transform*_) was used to calculate the optimal weight, 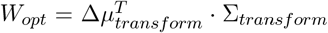, and the final estimated d’ was calculated by *W*_*opt*_ · *Δµ*_*test*_ and taking the square root of the result.

The technique from [23] was reimplemented in Python from their original Matlab code. Neural population responses from two conditions were used to estimate 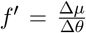, by taking the difference of their mean response normalized by the difference in stimulus. The covariance was estimated in a similar manner as well, except by taking the average of the two stimulus-specific covariances rather than normalizing by the difference in stimulus Δ*θ*. Linear fisher information was estimated from these quantities using partitioned matrix inversion techniques from [53]. To estimate the inferred information limit from the population response, a curve was fit via least-squares regression to the data 1*/I*_*fisher*_ vs 1/*N*, discarding the data from the first 10% of cells (so the curve starts fitting from 10% of the total cell population size, rather than from the first cell). This was done to provide a better linear fit in line with the theory laid out in the authors’ original work.

### Parameters used for the figures

*Figure 1*

The histograms of NCs in Fig. 1c,f,i were generated by simulating activity from a population of neurons with *N* = 1000. The neural activity followed von mises (see ‘Continuous Estimation’ in Simulations section above) tuning with *κ* = 0.75, and activity parameter *A* = 15. Neurons were presumed to have von mises tuning curves with preferred stimuli chosen uniformly along [0, 2*π*]. For the conditions where information limiting noise was present (*σ* > 0), the standard deviation of the input noise was set at 5 deg. When gain fluctuations were present in the simulations, *α* = 2. 5000 sample responses were generated from a uniform sampling of stimuli over the stimulus range [0, 2*π*]. The responses simulated neural activity within a time bin with width 120ms. For Fig. 1d,g,j, parameters for the simulations used to generate the scatter plots were identical to those specified above with the following differences: *κ* = 3, *A* = 10, and the number of iterations for each condition was 300. The decoding window for the decoder was 120ms. For Fig. 1e,h,k, the number of neurons along the x-axis was chosen with 20 steps with linearly spacing from 50 cells at the fewest to 2000 neurons at the largest population size. *κ* = 2, *A* = 8, *σ* = 5 deg. Neurons were presumed to have von mises tuning curves with preferred stimuli chosen uniformly along [0, 2*π*]. The number of iterations was 100. Stimuli was generated by uniformly sampling 5000 samples from U(0, 2*π*) and the tuning curves were used to estimate the continuous response to each stimuli, assuming a 120 ms decoding window. We verified that the results were insensitive to the choice of the model parameters and the differences were for display purposes only.

*Figure 2*

The number of neurons chosen for simulations were [100, 250, 500, 750, 1000], and the number of samples per condition were [25, 50, 100, 250, 500]. Values were chosen to cover an empirically feasible range while being able to display some level of asymptotic behavior. Neurons were presumed to have von mises tuning curves with preferred stimuli chosen uniformly along [0, 360] deg. The number of samples used for training the SVM was 75% of the total trials for the given iteration, of which there were 100 iterations per condition (cell count and sample count pair), with a 3-fold cross-validation procedure performed using the sklearn python package, selecting the best model while ranging over a SVC with parameters *max*_*iterations* ∈ [100, 1000], and regularization parameter C∈ [10^−4^, 10^2^]. The ground truth d’ values were [2,6,10], as labeled. For the matrix *D* described in the simulation section above, the 5 largest eigen-values were multiplied by 5 to increase the level of (non information-limiting) correlation in the population response, to provide more challenge to the analysis techniques used. For implementation of the method from Rumyantsev et al. [22], data was reduced to 5 PLS dimensions, consistent with their original report.

*Figure 3*

For each dataset, neuron counts were selected with 10 steps with geometric spacing from 50 to the maximum cell count for each session (generally in the range of ∼14,000-20,000 cells). Each decoding split was performed with 50 iterations. The *λ* parameter for the ridge regression based method used in [27] was set to 1.

*Figure 4*

Cells counts were selected along the range of [25, 50, 100, 250, Max]. Maximum recorded cell counts for the sessions ranged from 500-850 cells. The widths of the integration windows were chosen from 20ms to 300ms in steps of 20ms. The number of iterations performed for each condition (cell count, integration window width) was 100. The number of samples used for training the SVM was 75% of the total trials for the given iteration, with a 3-fold cross-validation procedure performed using the sklearn Python package, selecting the best model while ranging over an SVC with parameters *max*_*iterations* ∈ [100, 1000], and regularization parameter C∈ [10^−4^, 10^2^].

*Figure 5*

The number of neurons used in each iteration was selected with linear spacing from 10 to the maximum number of cells for the given session, with 10 steps in total. There were a total of 50 iterations performed for each cell population size in each experimental recording session. The parameters for the ZIG decoder [41], were *n*_*epochs* = 1800, *learning*_*rate* = 0.01, *gen*_*nodes* = 30, thresh = 0, *gam*_*shift* = 0.0001, *bin*_*len* = 181. The decoder was trained on 60% of the total data and the remaining 40% of data were split into half test and half validation.

## Code availability

The code used for generating all of the figures (main and supplementary) can be found at https://github.com/wei-bbc-lab/Split-Trial-Analysis.

## Acknowledgments

This research uses datasets from a number of previously published studies. We would like to thank the authors of these studies for generously sharing the data or making their data publicly available. We thank Thibaud Taillefumier, Zhongxuan Wu for helpful discussions, and Wilson S Geisler, Robbe Goris, Rong Zhu, Shizhe Chen for feedback on an earlier version of the paper. D.L. is supported by an GRFP fellowship from U.S. National Science Foundation. X.X.W. is supported by a Sloan Research Fellowship from the Alfred P. Sloan Foundation. This research is also supported by the NIH BRAIN Initiative and National Institute On Drug Abuse under Award Number R01DA060742. The content is solely the responsibility of the authors and does not necessarily represent the official views of the funding agencies.

## Supplementary Information

### A. Implementations of three alternative methods

#### A1. Standard (naive) decoding approach

According to the Cramer-Rao bound, the variance of an unbiased estimator approaches the limit of 1/*I*(*θ*) where *I*(*θ*) is the Fisher information contained in the population response about the variable *θ*. When we take the variance of our estimators’ decoding error as neuronal population sizes increase, we are essentially measuring the convergence to the information-limiting fisher information. Without using tailor-made tools and techniques, this is the most intuitive approach to estimate information limits in empirically recorded neural data, and so was chosen as our ‘naive’ approach for comparison purposes in this work. The idea of using a cross-validated decoder to estimate information-limiting noise was proposed in [5]. Results showing the performance of this method are shown in Fig. 2.

In a discrete classification task, we approximate the Fisher information instead using a measure of discriminability (i.e., d-prime), which can be viewed as 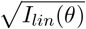, the square root of the linear Fisher information [5, 12]. The intuitive justification follows that of the continuous case otherwise. The results on the performance of this method are shown in Fig. S4.

Overall, the main problem with this standard decoding approach is that it exhibits a bias for over-estimating of the magnitude of the information-limiting noise. For the case of estimating d-prime, it generally under-estimates the ground truth d-prime values. Such estimation biases reduce as the number of neurons used increases.

#### A2. Method of linear extrapolation

The implementation of the technique from [23] was done by directly adapting the Matlab code (shared by the authors in their original report [23]) to a re-implementation in Python. In short, fisher information was calculated from population responses as neurons were added one at a time, and then a line was fit to the data when plotted 1/*N* vs 1/*I*(*θ*). The y-intercept of the line, when extrapolated, would give the *I*_∞_ estimate. In practice we chose to start projection from the first 10% of the population (so after the first 100-200 cells), for reasons to be described below.

The approach of [23] was essentially to note that after a sufficient amount of neurons are present in the population, the amount of information incremented by each additional neuron can be viewed as essentially constant, such that increasing population size leads to linear increases in information (as measured by fisher information). The intuition behind needing a “sufficient” amount of neurons before linearity can be observed primarily arises from the fact that one first needs enough neurons to have a complete (coarse) representation of the stimulus space. Once the stimulus space is properly spanned by the existing neurons’ tuning curves, then any stimulus can be represented by population activity and each additional neuron adds a constant amount of additional information.

A potential point of concern for the method is that the linear projection to the *I*_∞_ value may be unstable depending on the size of the population and the magnitude of the (information-limiting) noise. If the population is too small, then additional neurons grant non-constant increases in population fisher information as they potentially cover unique portions of stimulus space. If the limiting noise is quite small, then the linear extrapolation can have difficulties getting the precise value because the fit is performed in 1/*I*(*θ*), and when *I*_∞_ is large, the ensuing fraction can be quite small and numerical instability may occur. In our simulations, we find that this method based on extrapolation [23] is numerically unstable and it depends on the number of trials and the number of neurons. Simulation results to demonstrate these points are shown in Fig. 2, Fig. S4, and Figs. S5-S7.

#### A3. Method of dimensionality reduction

The approach from [22] for classification between two stimuli was implemented as described in the methods section of their own report. In essence, the neural response data is split into 3 partitions, which we’ll denote *X*_*train*_, *X*_*weight*_, *X*_*test*_ which are used for learning the transformation, obtaining the optimal weight, and calculating the final d’ value, respectively. The first partition *X*_*PLS*_ is used to train a partial least-squares regression model, which reduces data down to a lower dimension. The authors discovered n=5 was optimal for their data, which we followed in our implementation as well. Then the model was used to transform another data partition, *X*_*train*_ into the low dimensional space. The resulting low dimensional data was used to estimate Σ = 0.5 (Σ_1_ + Σ_2_), the low dimensional noise covariance matrix, where Σ_*i*_ were the noise covariance matrices for the dimension reduced neural responses from either of the two stimulus conditions. The optimal transformation weight was then found calculating *w*_*opt*_ = Σ^−1^ · Δ*µ*_*weight*_. The final partition is used to calculate d’ by performing 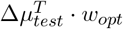.

In general, the method operates by assuming that the main drivers of neuronal noise can be captured in a low dimensional subspace, and by reducing the dimensionality, the noise covariance matrix estimation is much more numerically feasible. It is in particular effective considering theories that information-limiting noise is restricted to a single direction tangent to the signal manifold [5]. However, it should be noted that the method, as described, only seeks to estimate the information for a finite number of neurons, namely, the neurons present in the population. Specifically, the method does *not* seek to estimate *I*_∞_. As a result, its primary use seems to be to best estimate the amount of information encoded by an empirically recorded neural population. As a result, the method is not optimal for detecting information-limits in a population, and is best used to estimate the maximal information that can be extracted from a given finite population. In practice, this allows the method to perform better than naive estimation procedures, but falls short of estimating the information-limiting noise in general, in particular when the information-limiting noise is small (Fig. 2, Fig. S4).

#### B. Model justification when only information-limiting noise and private neural noise is present

In this section, we provide a justification for the split-trial analysis in the context of a standard neural population model in which only information-limiting noise causes noise correlations.

##### Model setting

Assume that a one-dimensional stimulus variable *θ* is represented by a neural population code with *N* neurons (*N* may approach infinity). Denote the tuning curve of the *i*_*th*_ neuron as *f*_*i*_(*θ*), which is a smooth function. *f*_*i*_(*θ*) does not need to be Gaussian function. We will assume that neurons exhibit Poisson spiking noise, although the arguments below may also work for other types of noise (such as negative binomial noise or Gaussian noise). Thus, the code can be expressed as the following:

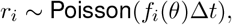

where *r*_*i*_ represents the activity of the *i*_*th*_ neuron.

We model information-limiting noise by injecting Gaussian noise to the stimulus

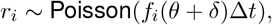

where the random variable *δ* denotes information-limiting noise and is shared by all neurons. Note that, in this model, noise correlations are only induced by information-limiting noise. The Poisson spiking noise is private to individual neurons. Although this setting is likely simplified, as in practice there may exist other types of shared neural variability that are not information-limiting, our simulation results in Fig. 1 show that it is already challenging to use standard decoding methods to estimate information-limiting noise.

##### Analysis of the split-trial approach

Suppose that we collect neural responses from this population of neurons (N neurons), and perform split-trial analysis. Our goal is recover the variance of information-limiting noise *δ*. When performing split-trial analysis, we split the neural population into two halves and perform Bayesian decoding on each split to obtain the Bayesian estimator of the stimulus. In performing Bayesian decoding, we simply assume that neurons exhibit independent neural noise.

Assume that information-limiting noise *δ* is small and the tuning curve is smooth, given a sufficient amount of data, the tuning curves for individual neurons can be accurately estimated. For a sufficiently large neural population, the Bayesian estimator based on each split, 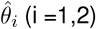, is expected to be approximately Gaussian distributed [54], centered at *θ* + *δ*. The reason is because Bayesian decoding in this case is essentially decoding *θ* + *δ* from a large neural population with independent neural noise. Thus, we can write

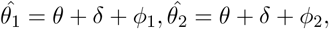

where *ϕ*_1_ and *ϕ*_2_ represent the decoding error of *θ* + *δ* for the two splits. Furthermore, because that neural noise is independent in the neural population (when treating the decoding problem as one that decodes *θ*+*δ*), the decoding error from the two splits should be independent. Therefore, under the above assumptions, *ϕ*_1_ and *ϕ*_2_ are independent random variables. Given these, a straightforward calculation (as shown in the Methods section) shows that the covariance of 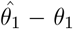 and 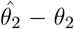 can be used to estimate the variance of information-limiting noise *δ*.

Suppose that the variance of information-limiting noise is *σ*^2^. As the variances of *ϕ*_1_ and *ϕ*_2_ should be nearly identical when *N* is large, for simplicity we will assume that they are both 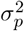. When using all *N* neurons to perform Bayesian decoding analysis, the variance of Bayesian estimator should be of the order of 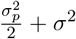. This is because integrating information from the two splits should reduce the variance of the estimator from the private noise by a factor 2, while preserving the variance due to the shared noise *δ*. Thus, using Bayesian decoding to estimate the variance of information-limiting noise suffers the problem of over-estimation, with an upward bias of 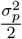. While this bias will converge to 0 when the number of neurons is infinite, practically the bias can be substantial, as we showed in our simulation results. Increasing the number of trials will not eliminate this bias.

For split-trial analysis, empirically the covariance estimator 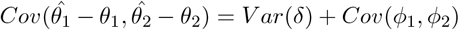 may exhibit statistical fluctuations, due to noisy empirical estimates of *Cov*(*ϕ*_1_, *ϕ*_2_) (see Fig. 1). However, the estimator should generally be unbiased as *Cov*(*ϕ*_1_, *ϕ*_2_) is expected to be 0. Our simulation results in Fig. 1 show that when the number of neurons becomes larger, the variability of this estimator reduces and becomes more accurate. For finite samples, the estimator is off by *Cov*(*ϕ*_1_, *ϕ*_2_). Practically, this amounts to 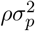, where *ρ* is the Pearson correlation of the samples of *ϕ*_1_, *ϕ*_2_. From Fig. S1, we see that *ρ* is generally small, with most values between [-0.1, 0.1]. For the sake of argument, suppose that *ρ* is 0.1, the error in estimating the information limiting noise should be about a fifth of the error of the standard (naive) method based on Bayesian decoding. Furthermore, our estimator does not suffer from the systematic overestimation of the naive method.

##### Remarks

The analysis above assumes that all noise correlations are caused by information-limiting noise. When other sources of shared neural variablity other than the information-limiting noise are present, one can still perform Bayesian decoding on the two splits. When the number of neurons is large, one may still approximately analyze the Bayesian estimator using a similar approach as above, i.e., 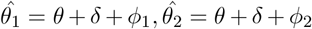. However, for a finite neuron number *N, ϕ*_1_ and *ϕ*_2_ may be correlated depending on the exact structure of the shared, non-information-limiting noise in the neural population. When *N* is large, *Cov*(*ϕ*_1_, *ϕ*_2_) should approach 0, otherwise there would be non-zero information-limiting noise shared by *ϕ*_1_, *ϕ*_2_ (which would contradict to model assumptions). Such a non-zero correlation between *ϕ*_1_ and *ϕ*_2_ may increase the error of our split-trial estimator. However, practically, we find that, at least for the case of gain fluctuations, our split-trial estimator still works well even when we use a decoder that assumes independent neural noise (see Fig. 1 and Fig. S2). In the next section, we perform additional analysis based on general Gaussian noise correlation structure, in the context of a neural population coding model assuming Gaussian tuning curves. These analyses provide additional justification for why we expect the split-trial analysis to work well in the more general setting.

### C. Model justification under more general assumptions on noise covariance

In this section, we will investigate the properties of split-trial analysis under more general noise assumptions. We will do so in the context of neural populations with Gaussian tuning curves. We let 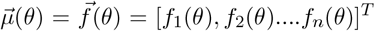. We will, for each stimulus parameter *θ*, let the inverse covariance matrix Σ^−1^(*θ*) be a diagonal matrix, diag 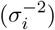, reflecting the fact that decoding of empirical data often assumes independence in activity across the population. Then population responses 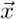 are drawn according to 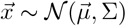, omitting *µ* and Σ’s dependence on *θ*.

#### C.1 Properties of the locally optimal linear estimator

First, we will investigate the mean and variance in the error of the locally optimal linear estimator (LOLE) [55, 5]. We will use these results for further calculations and for assessment of the split-trial technique. Note that by using the locally optimal linear estimator, we are assuming that the decoder has knowledge of the noise correlation structure that is induced by non-information-limiting noise. The locally optimal linear estimator for 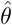 can be written as:

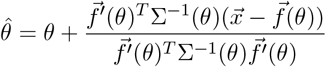

known to be optimal when estimating a parameter value near *θ* [55, 5]. Starting with the denominator of the fraction:

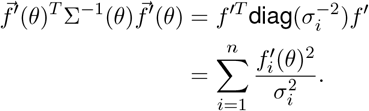

We evaluate the numerator similarly:

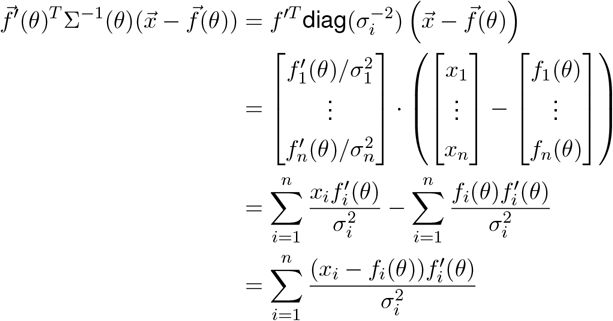

We can then combine terms and evaluate the decoding error:

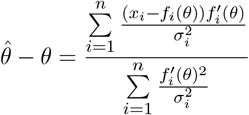

From which we can see that, were we to take the expectation of 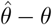, we would find the error has mean 0 since E[*x*_*i*_] = *f*_*i*_(*θ*).

We proceed with the variance calculation:

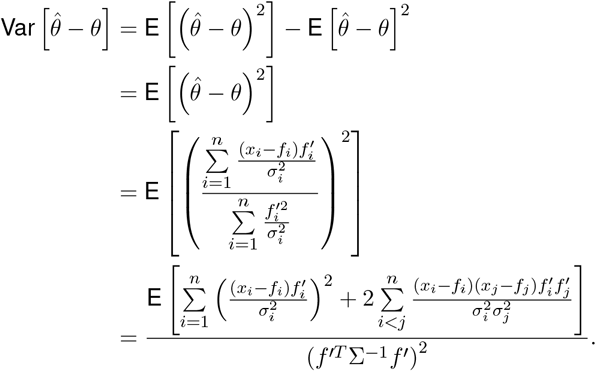

We’ll evaluate each term individually starting with the diagonal term in the numerator:

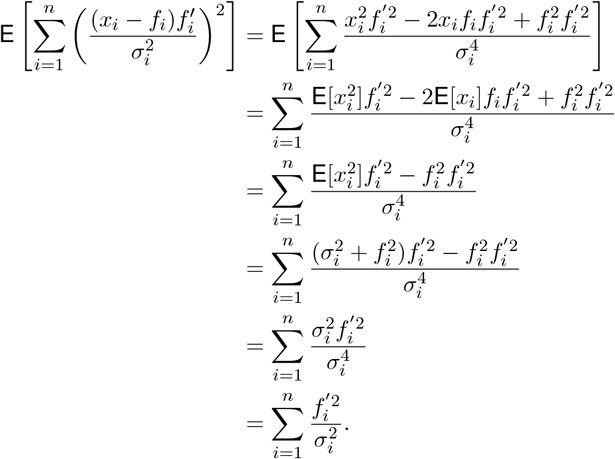

Now we address the numerator cross-term:

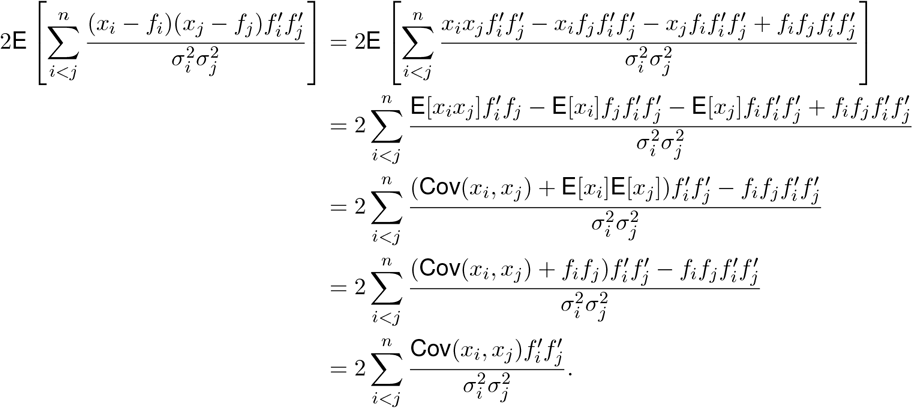

So we have that

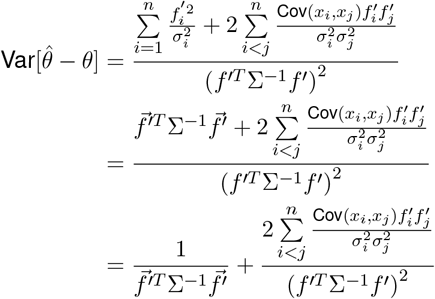

The denominator of this expression is the linear Fisher information arising if the population exhibits a covariance that aligns with the assumption of the estimator (which in this case was that the activity is independent), whereas the numerator contains the Cov(*x*_*i*_, *x*_*j*_) term which would track the true covariance values. If the estimator has perfect knowledge of the ground truth then 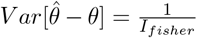 and the Cramer-Rao bound is achieved as Cov(*x*_*i*_, *x*_*j*_) = 0, ∀*i*≠ *j*. If there is a mismatch in ground truth and presumed covariance structure, then the variance is larger by the second term in the above equation. The second term cannot be negative as that would violate the Cramer-Rao bound. For the intuition behind this, consider: if Cov(*x*_*i*_, *x*_*j*_) < 0, 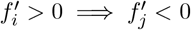, and the numerator of the sum remains positive. All other terms are squares and forced positive. In other words, if two variables are negatively correlated, then one must increase firing rate when the other decreases, thus the product of the *f*^*′*^ terms is negative whenever Cov(*x*_*i*_, *x*_*j*_) is negative, ensuring positivity of the second term, the case where variables are positively correlated follows.

#### C2. Covariance calculations

In this section, we will examine the performance of split-trial analysis under different noise regimes, characterized by assumptions on the ground truth noise covariance matrices, Σ. Let 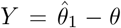 and 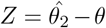 be the error estimate from the first and second population splits given stimulus *θ*, respectively. Additionally, let each tuning curve be defined by mean *g*_*i*_, *h*_*j*_ with variance *η*^2^, *ν*^2^, respectively as well. We will use only a two-fold split for proceeding calculations. This is equivalent to considering Σ as a block matrix 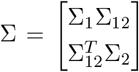 where e.g. *Y* is calculated as above using Σ_1_. Specifically, we would represent 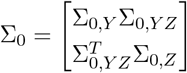. Additionally, we will presume that our covariance matrix has the general structure of Σ = Σ_0_ +*ϵf*^*′*^*f*^*′T*^, where *ϵ* is the magnitude of the information limiting noise, and Σ_0_ is a covariance matrix with no information-limiting components, following the convention of [5].

##### C2.1. Split-trial analysis when only information-limiting noise is present

We will start with special case for which 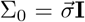, a diagonal matrix. In this case, all noise correlations are caused by information-limiting noise. Recall 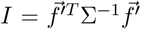 is the linear Fisher information for a population with tuning curves 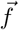 and covariance Σ.

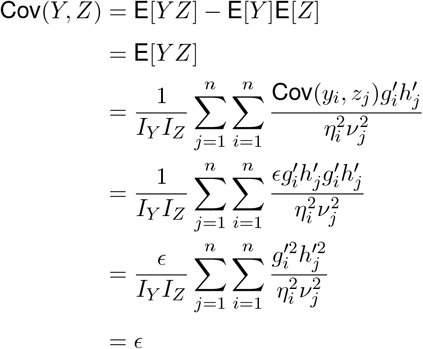

which confirms the validity of our estimator in the case where information-limiting noise is added to an otherwise independent population, in the case of a population of neurons with Gaussian tuning curve and Gaussian noise. These results complementary to the results reported in Section B.

##### C2.2 Split-trial Analysis with general noise covariance matrix

In this section, we will relax assumptions on Σ_0_ and examine the general case. Specially, we allow other forms of noise correlations that are not caused by information-limiting noise. Let 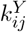 be the element of 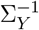 in the *i*th row and *j*th column and *C*_0,*ij*_ be the element in the *i*th row and *j*th column of Σ_0_.

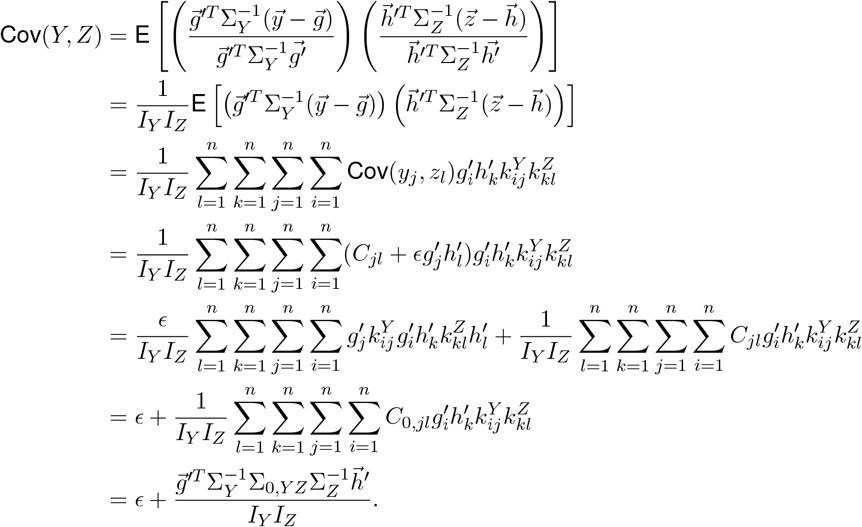

We examine this second term causing deviations away from *ϵ* in the following section. Although, one can note that, if the only noise present in the recorded population is from information limiting noise (i.e. the covariance Σ_0_ is diagonal), then Σ_0,*YZ*_ is the zero matrix, and the second term vanishes, consistent with previous calculations.

##### C.2.3 Examination of the error term in split-trial analysis with general noise covariance matrix

Here we attempt to develop an understanding of the second term in the above calculations for Cov 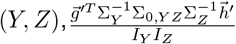. Specifically we will be focusing on the numerator for this expression: 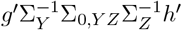. It will be convenient to work with block matrices for this section, so we will be representing Σ_0_ as 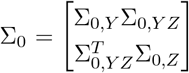. We will also be representing these tuning curves in a block format: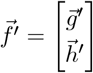.

###### Discussion on estimator knowledge of covariance structure

In this section we are going to examine an alternate form to our error term and from which derive some useful properties of the split-trial analysis. We will begin with establishing some quick identities for linear fisher information, using *I*_*Y*_ as an example. Proceeding from the assumption that Σ_*Y*_ = Σ_0,*Y*_ + *ϵg*^*′*^*g*^*′*^*T*, we can use the Sherman-Morrison formula to obtain:

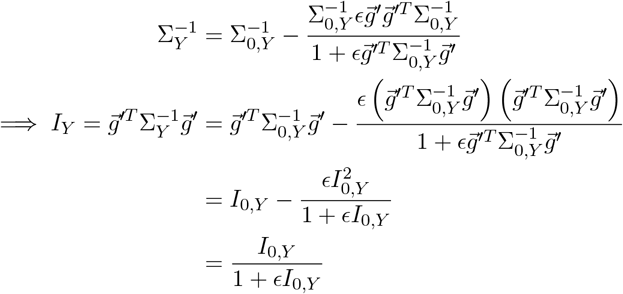

as was similarly presented in [5]. We can now return to our error term with these calculations in mind. Specifically, we can say:

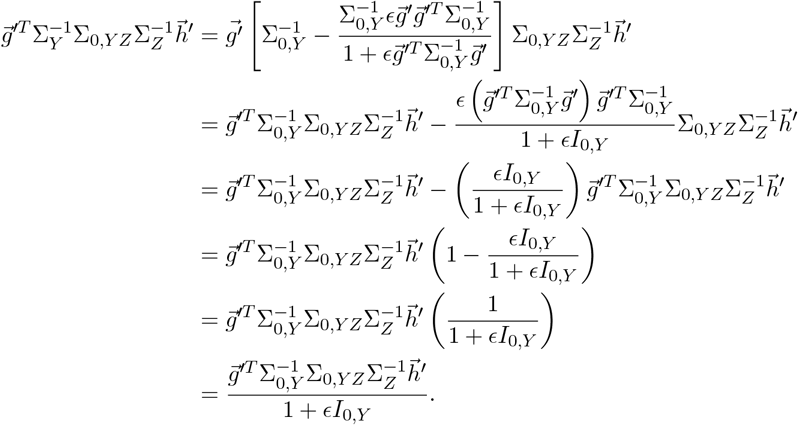

A similar argument using 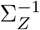 gives us that

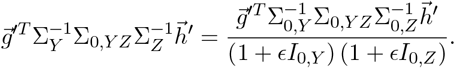

Now we can use the previously established substitutions for *I*_*Y*_, *I*_*Z*_ established above to write

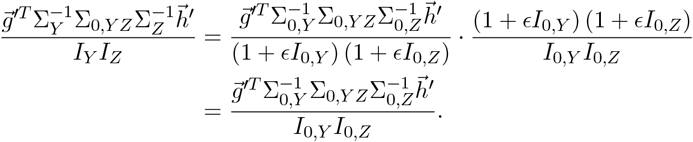

What this rewriting shows us is that the split-trial analysis error term is invariant to the decoders’ knowledge of the covariance structure of the data. The newly acquired term we have would arise if, in the original calculation, all the covariance terms (from, say, 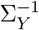) were replaced with the same terms in the analogous non-information limited matrix 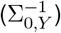. Indeed, this also demonstrates that the error in the split-trial estimation procedure is entirely dependent on the properties of the non-information limited noise in the system, which will be exploited in future sections as we will primarily perform calculations using this non-information limited form of error.

###### Establishing bounds on recovered information-limiting noise values

Limiting the influence of the above mentioned error term is necessary in order for the split population analysis to return the ground truth *ϵ* noise, and in this section we shall endeavor to establish a bound on this error. *Assumption 1*: we will need to make the assumption that the recorded neural population has some base level private neuronal noise, *σ*_*min*_. This guarantees that all of the covariance matrices we work with are strictly positive definite (rather than merely semidefinite), which ensures that all of the quadratic forms encountered are strictly positive (e.g. 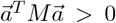), and that all eigenvalues are also strictly positive. *Assumption 2* will be to also carry on the assumption from the previous section that the subpopulation splits have identical statistical properties such that *I*_0,*Y*_ = *I*_0,*Z*_ = *I*_*half*_.

We will primarily be examining the numerator of the error term, 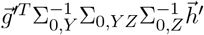. It is at this juncture that we wish to point out the remarkable similarity our error term 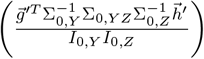 has to the so-called “canonical correlation”: 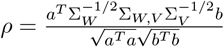. The similarities in the two expressions are in some sense unsurprising as canonical correlation analysis (CCA) is interested in assessing the correlations between two distinct datasets, a scenario that our split-trial method constructs by design. This connection and its repercussions will be explored further in the next section. For now, we shall primarily focus on establishing a finite upper bound to our error term.

We first will rewrite 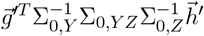 as 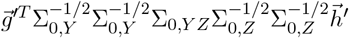. We can then obtain

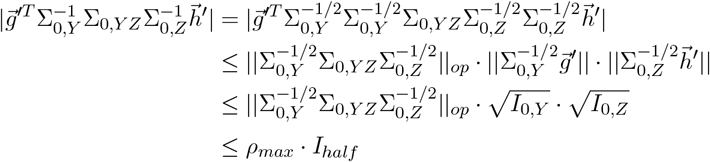

where the second line is an application of Cauchy-Schwarz,|| ||_*op*_ is the operator norm, and *ρ*_*max*_ is the maximum canonical correlation between subpopulations *Y* and *Z*. The equivalence between the canonical correlations and the operator norm in the last line comes from established results on CCA and singular values as shown in [56]. From here we can establish 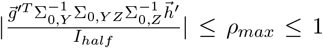 and we conclude 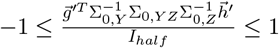.

As one can perform arbitrary partitions of our neural population, it is reasonable to consider 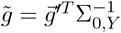 and 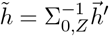to be independent random vectors with respect to one another. We assume no structured correlations in the network and additionally that Σ_0,*YZ*_ is independent of 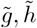. Given that, we can consider 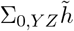 and 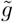 random vectors with respect to each other. It is known [57] that the angle between two random vectors Θ is orthogonal with scaling 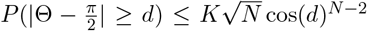 for 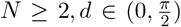, where N is the number of neurons in our total population, and K is a fixed constant. This means that there should be an exponential scaling (towards 0) in the numerator of our error term. This exponential scaling causes a rapid convergence of the inferred noise in split-trial analysis towards ground truth *ϵ*. The convergence rate in this regime is faster than the convergence of the standard method based on direct decoding and the Cramer-Rao bound.

In the worst case however, we can consider Σ_0,*YZ*_ to have values that are dependent on 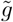 and 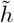. Consider a special example that the neural population exhibit population-wide structured correlations of a particular form such that 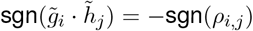, where sgn is the sign operator. Assume that these structured correlations have magnitude *ρ*_*gain*_, in such a case we would have 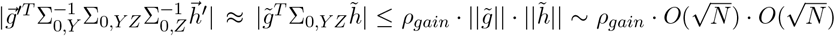, since 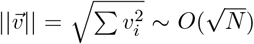. Our estimator would therefore return values of the form 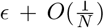, in line with the standard approach’s scaling in this special case. Our simulation results suggest however, that even in one of these worst cases (structured multiplicative gain fluctuations with information limiting noise), split-trial analysis still performs better than the standard approach (see Fig. S2).

**Figure S1:**
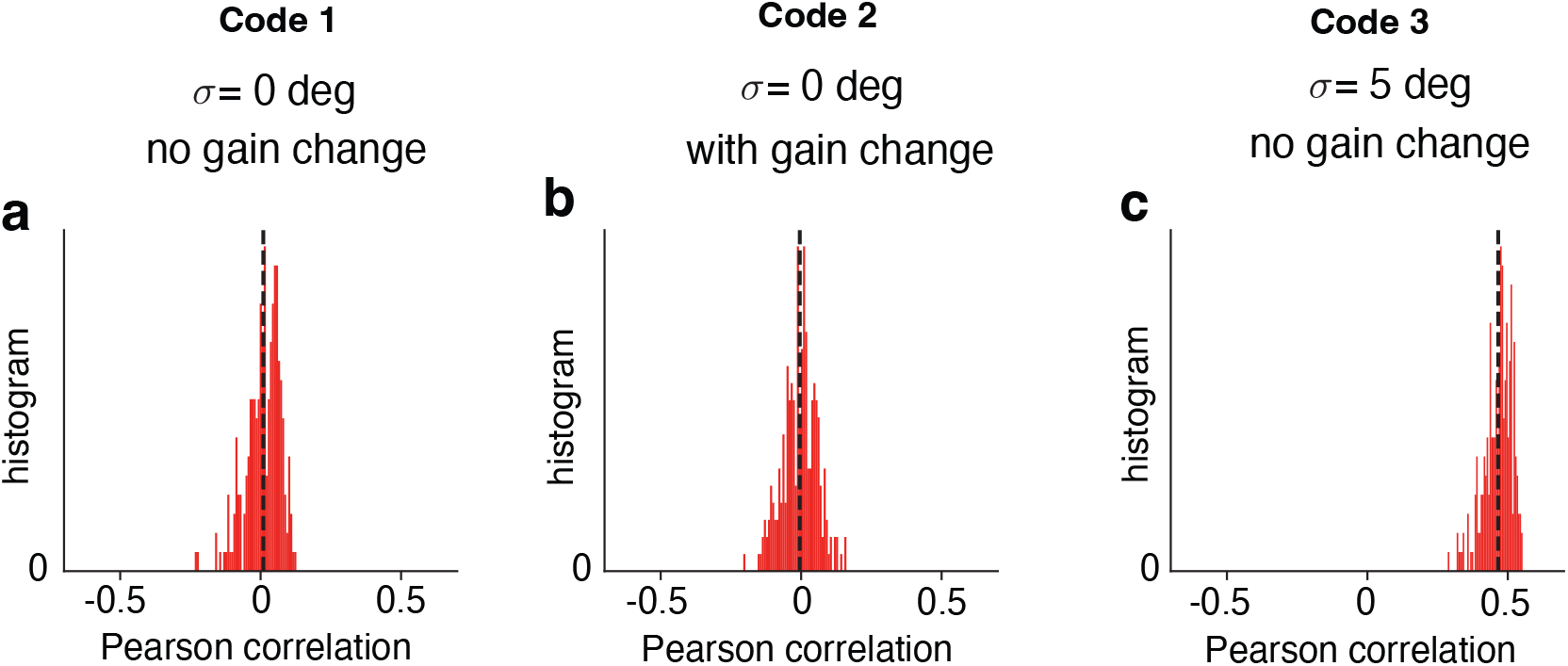
Histograms of the Pearson correlation values between the two splits for the three neural population codes studied in Fig 1. For each code, we perform split-trial analysis by generating simulated response, decoding the two splits and calculating the correlation between the decoders from the two splits. The results show that when information-limiting noise is 0, correlation values are generally around 0. In the presence of information-limiting noise, correlation values are generally positive.

**Figure S2:**
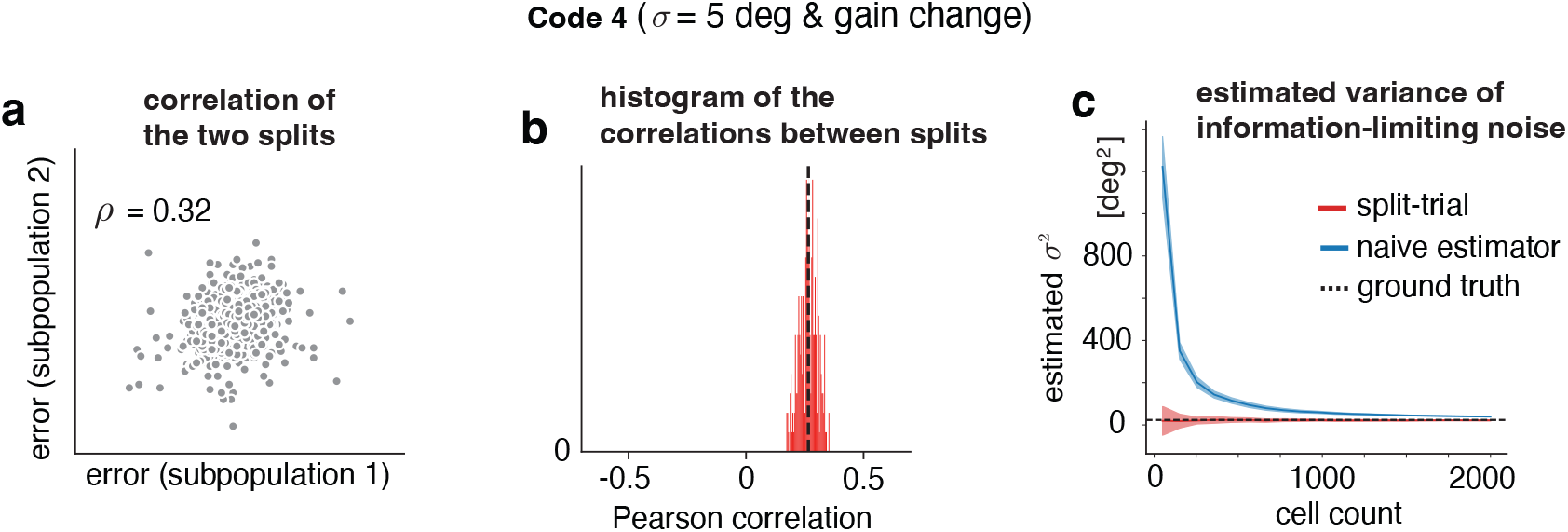
Split-trial analysis for another neural population code. We further constructed a fourth neural population code (Code 4) by incorporating both information-limiting noise (*σ* = 5 deg) and shared gain fluctuations (non-information limiting). We find that split-trial analysis can efficiently recover the information-limiting noise in the presence of both information-limiting and non-information limiting noise.

**Figure S3:**
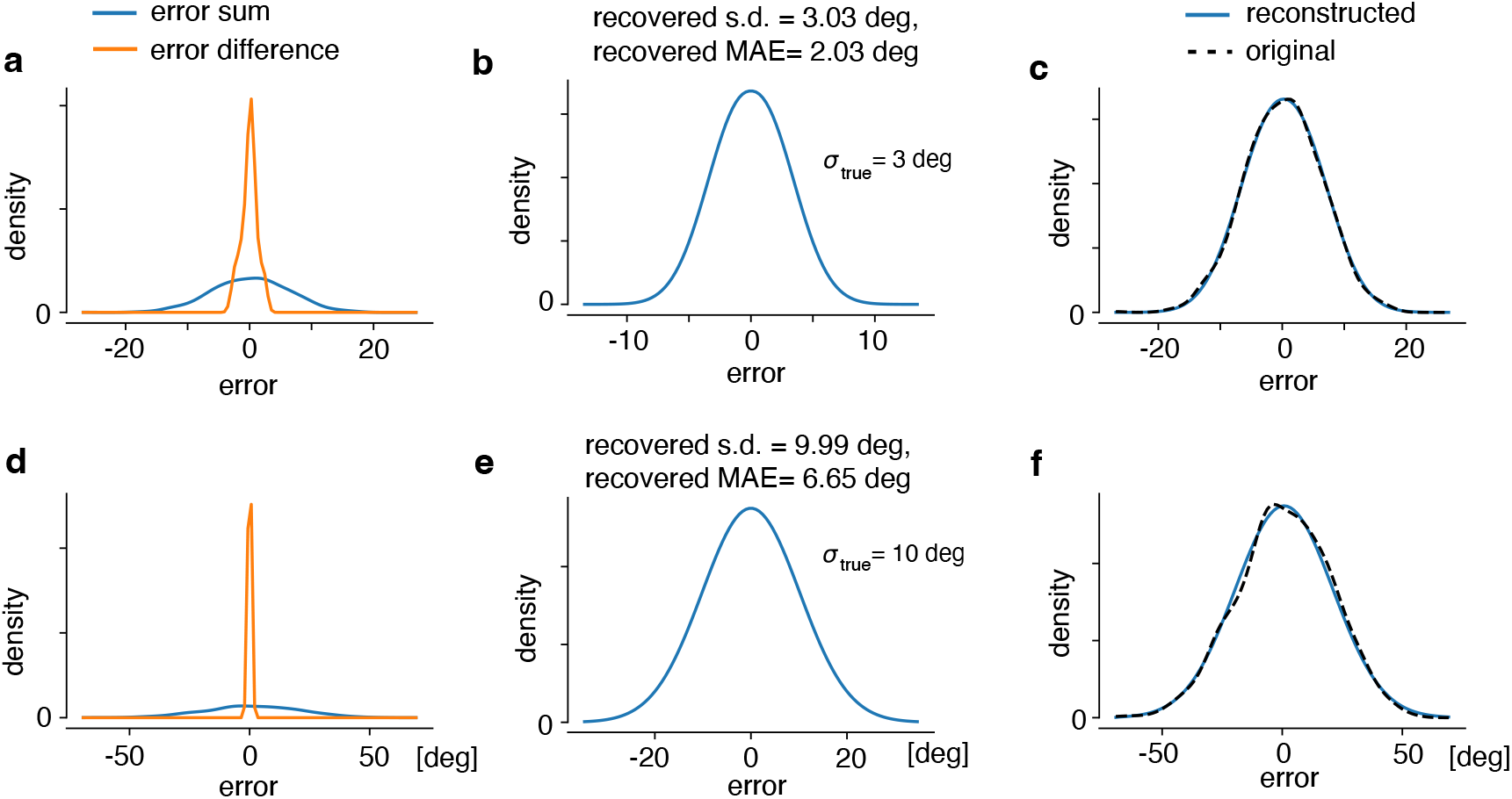
Validation of deconvolution based approach for estimating information-limiting noise. Data is simulated in similar schemes as **Fig. 3**, but the information limiting *σ* is estimated using a deconvolution approach.*Top row:* data generated with a ground truth information-limiting *σ* = 3, *bottom row:* data generated with a ground truth information-limiting *σ* = 10. **(a**,**d)** The sum and difference of errors across both splits. **(b**,**e)** The estimated error distributions for the two datasets, with the estimated *σ* and MAE listed. **(c**,**f)** Using the deconvolving filter estimating the error distribution, we can reconstruct the original error sum distribution (solid blue) and compare it against the actual original distribution (dashed).

**Figure S4:**
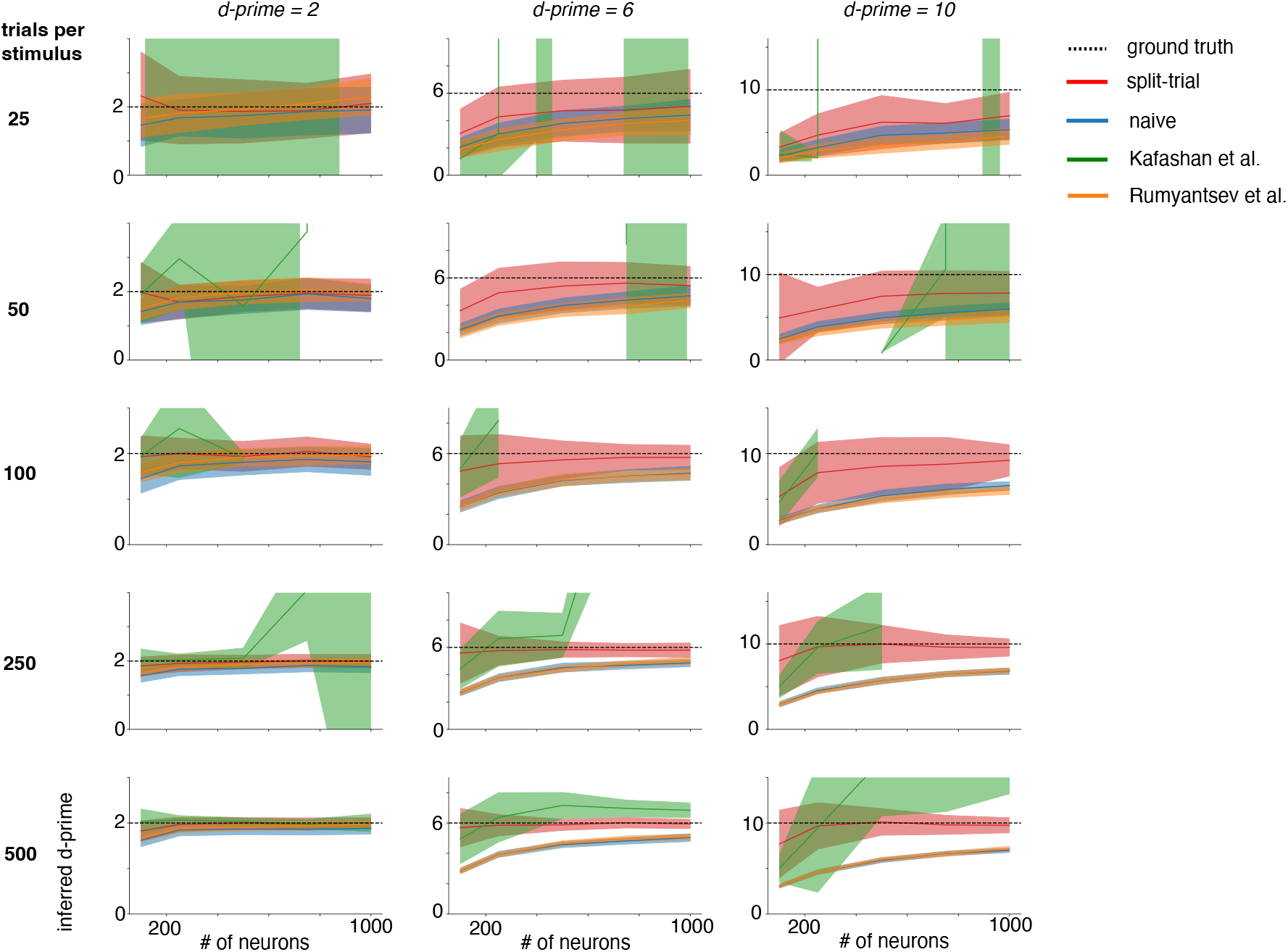
Benchmarking the performance of split-trial analysis and three alternative methods on inferred the ground truth d-prime (related to Fig. 2). This figure shows the results for all sample/stim and cell population combinations used (related to Fig. 2). Note that the portions of the curves for which the method from [23] disappears due to a preponderance of unbounded information estimates.

**Figure S5:**
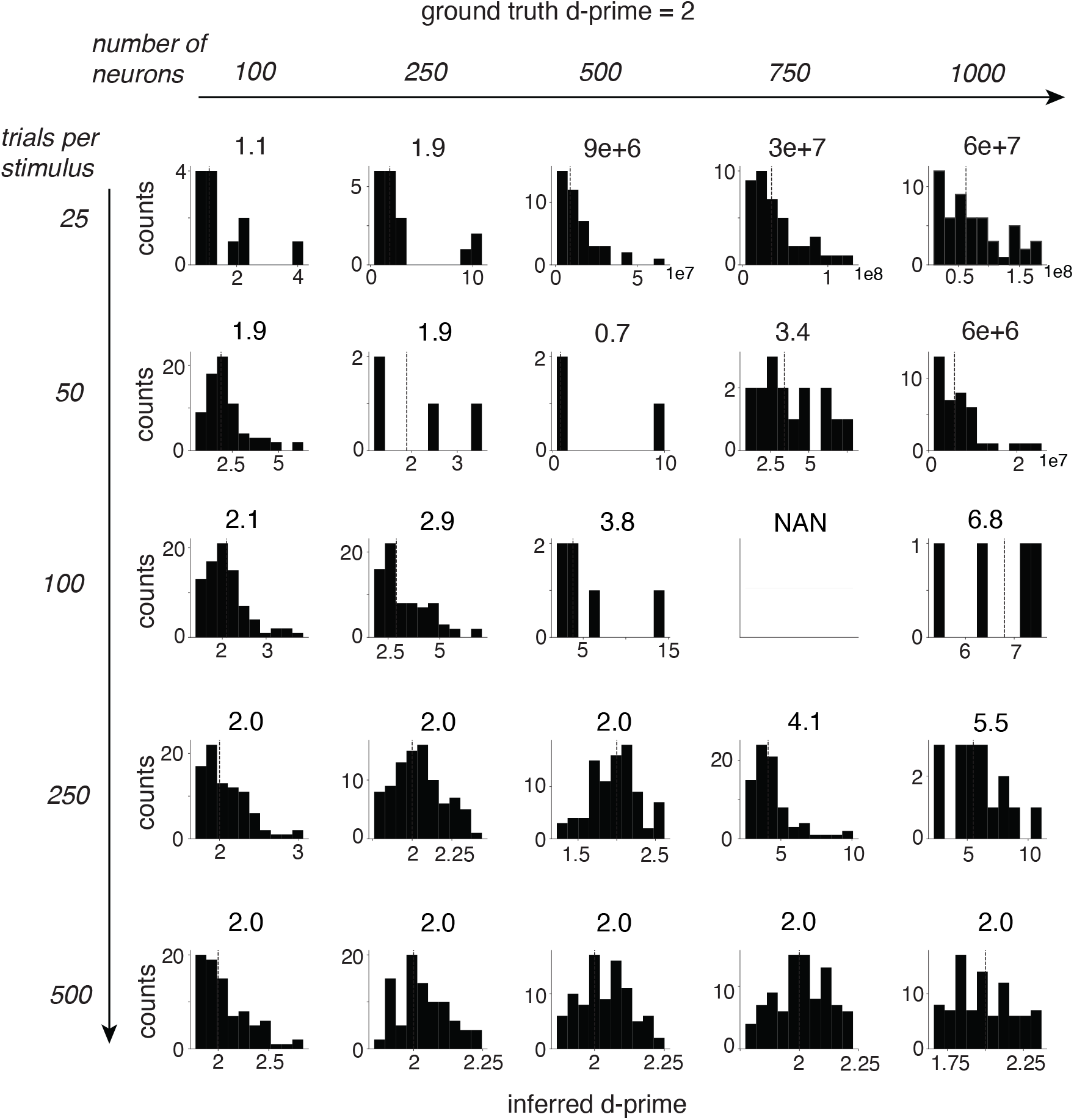
Histograms of estimated d-prime limit for the method from [23] based on extrapolation. The ground truth d-prime in this case is 2. For each panel, the median of the estimated d-prime values across 100 simulations was shown on the top. Note that, when there are few samples, the linear extrapolation method is highly unstable. Additionally, certain combinations of large cell count and large samples may be biased towards classifying a population as having unbounded information. This is likely due to the particular bias-corrected fisher information calculation developed used in the original report. When the estimated information is unbounded, such estimates were not shown in the plots (thus the sum of the counts in the histogram may be smaller than 100). In some case, none of the 100 simulations results in a finite estimate of the d-prime values, resulting in all NANs.

**Figure S6:**
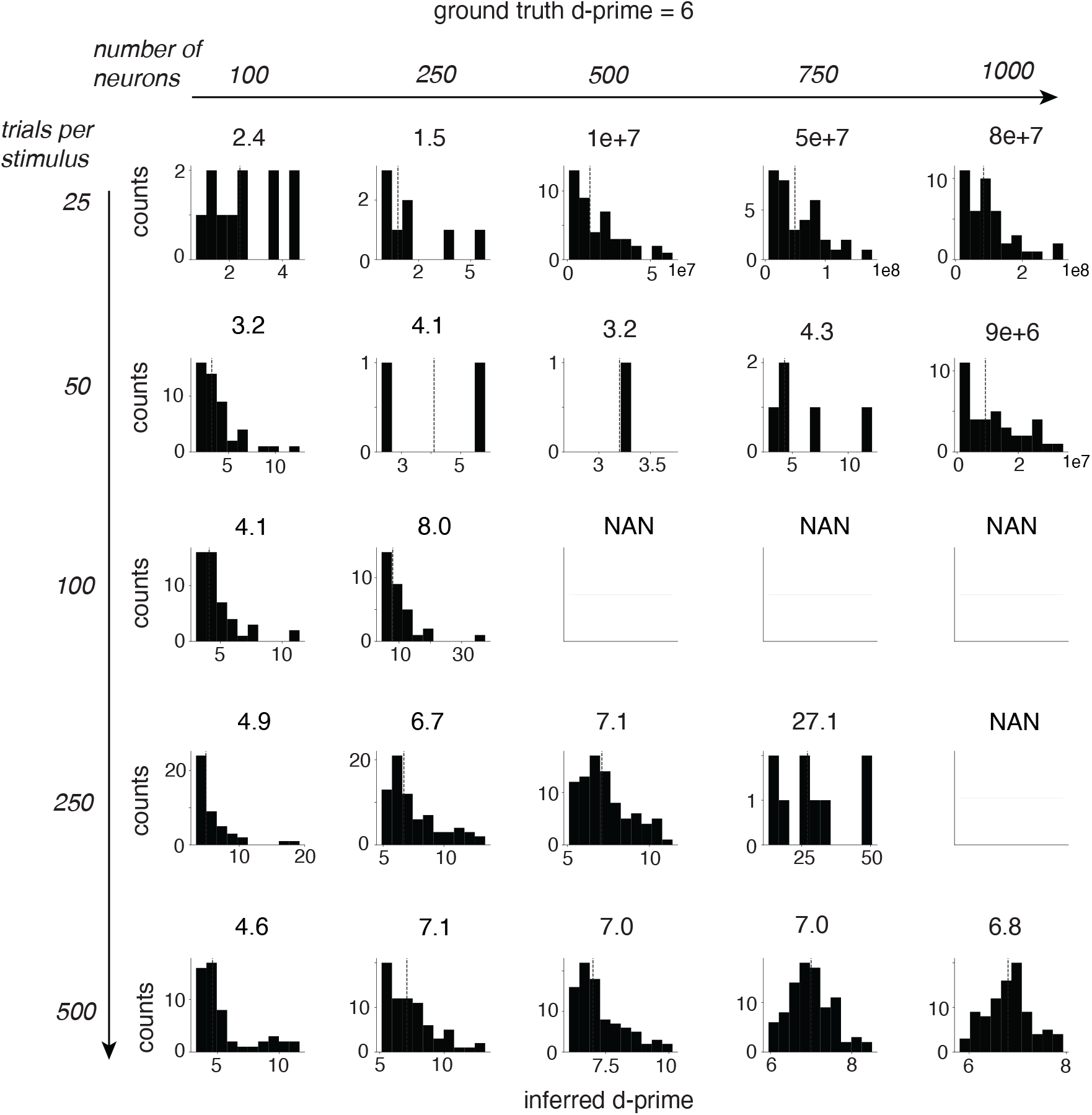
Histograms of estimated d-prime limit for the method from [23]. The ground truth d-prime in this case is 6. Same convention as Fig. S5.

**Figure S7:**
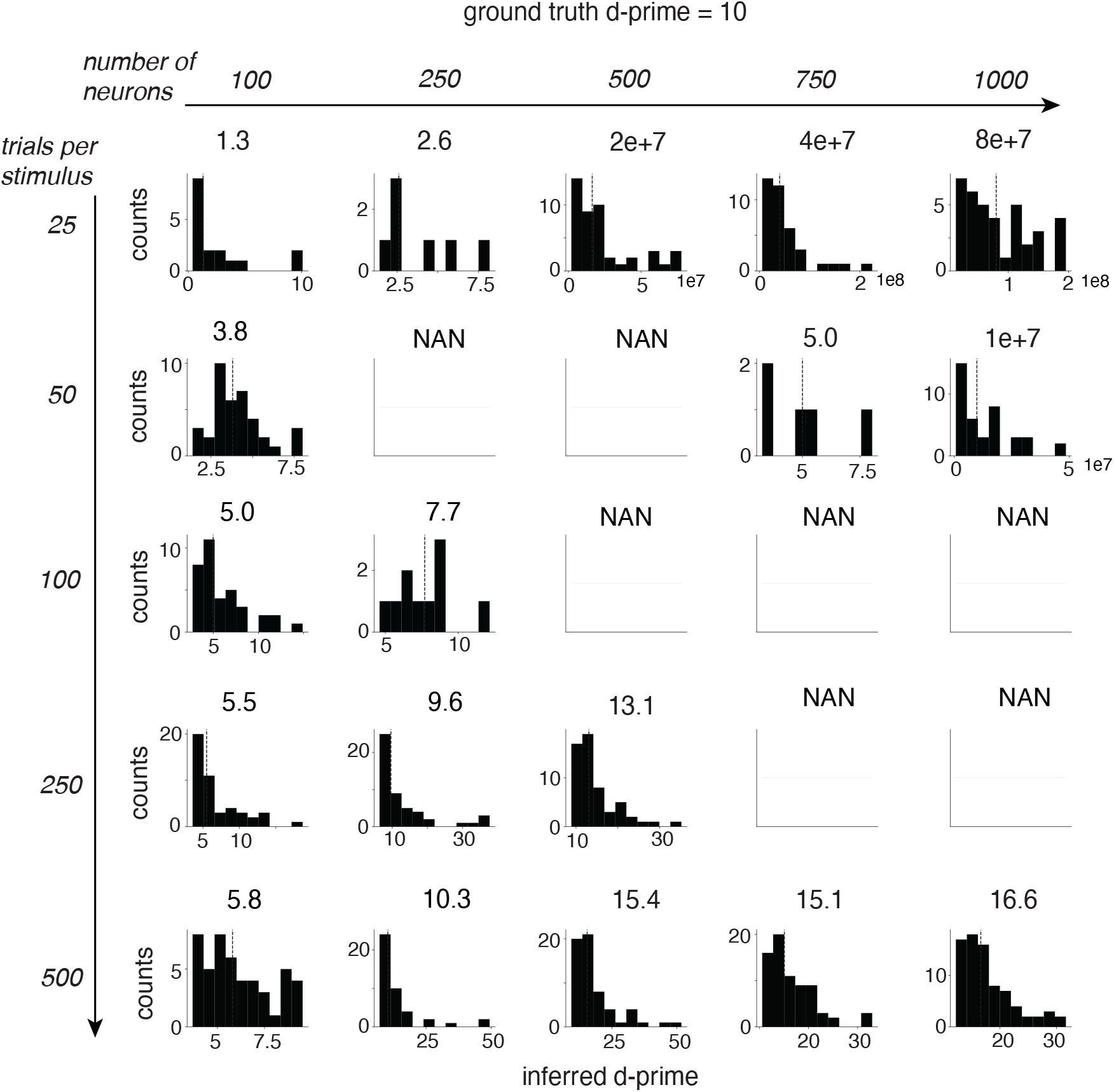
Histograms of estimated d-prime limit for the method from [23]. The ground truth d-prime in this case is 10. Same convention as Fig. S5.

**Figure S8:**
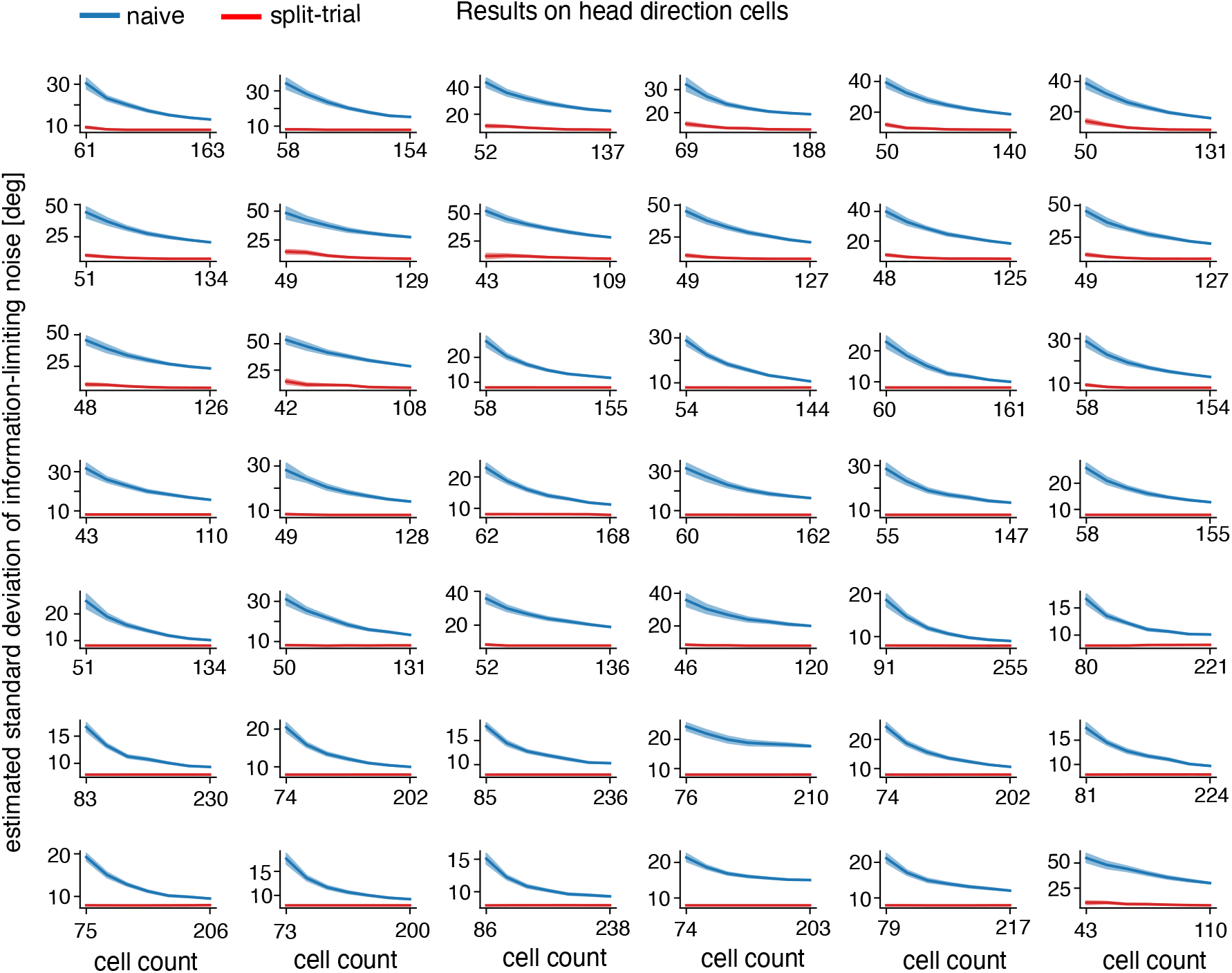
Individual session estimates for the information-limiting *σ* from [34] for both split and naive approaches. Plots were started from intermediate population sizes for visualization purposes.

**Figure S9:**
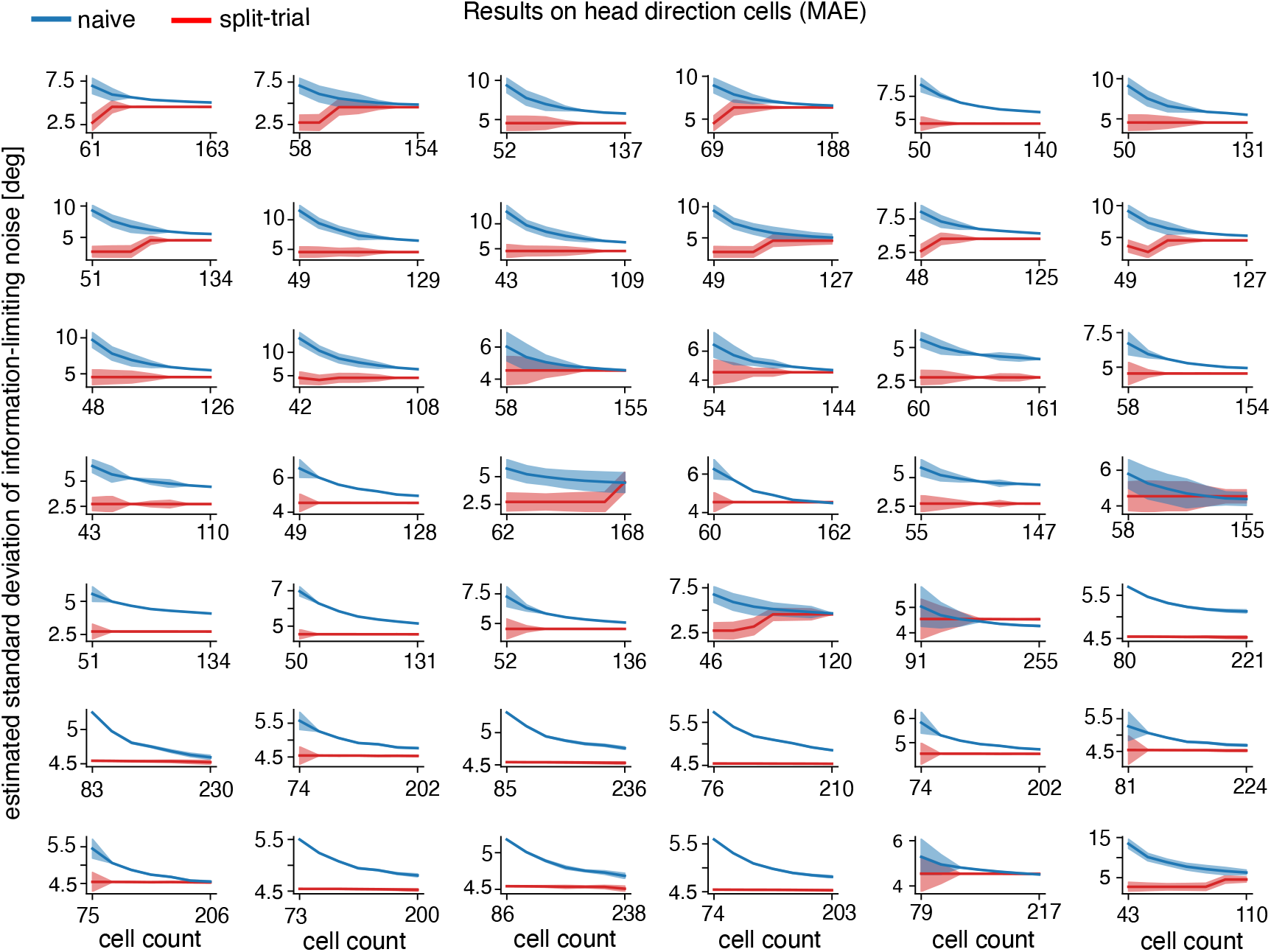
Individual session estimates for the median absolute error (MAE) of information-limiting niose from [34]. Red: results based on split-trial analysis based on the deconvolution approach. Blue: results based on direct decoding using a ZIG Bayesian decoder[41]. Plots were started from intermediate population sizes for visualization purposes.

**Figure S10:**
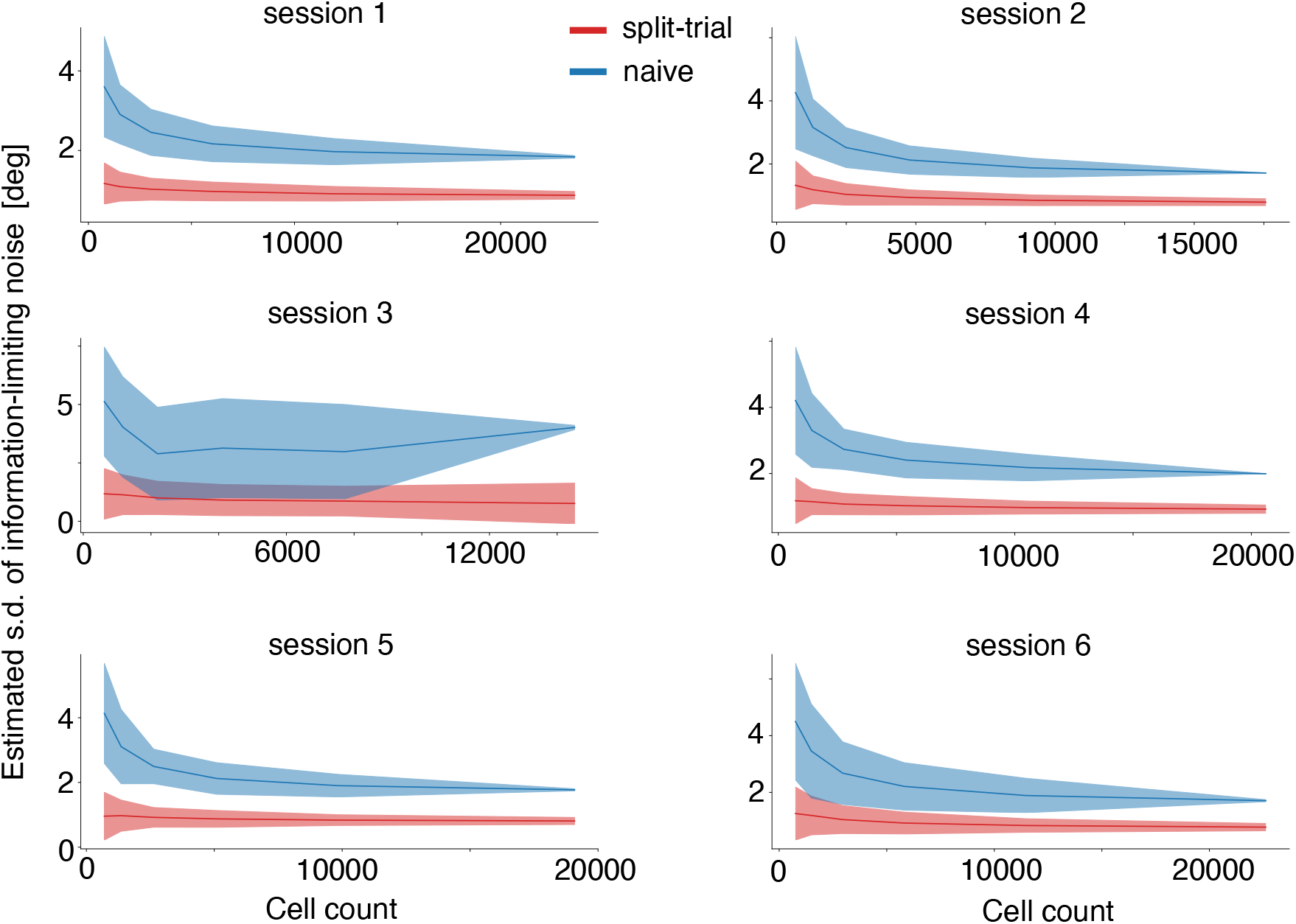
Results on the inferred standard deviation of information-limiting noise from the uniformly sampled orientation condition. Results from 6 individual sessions were shown. Red: estimates from split-trial analysis. Blue: estimates from the direct (naive) decoding analysis. We observed that the naive estimate of *σ* for session 3 increases slightly after ∼ 10,000 cells. The odd level of variability in this session made display challenging so was excluded from the visualization of Fig. 4c. Error bars are standard deviation over all iterations.

**Figure S11:**
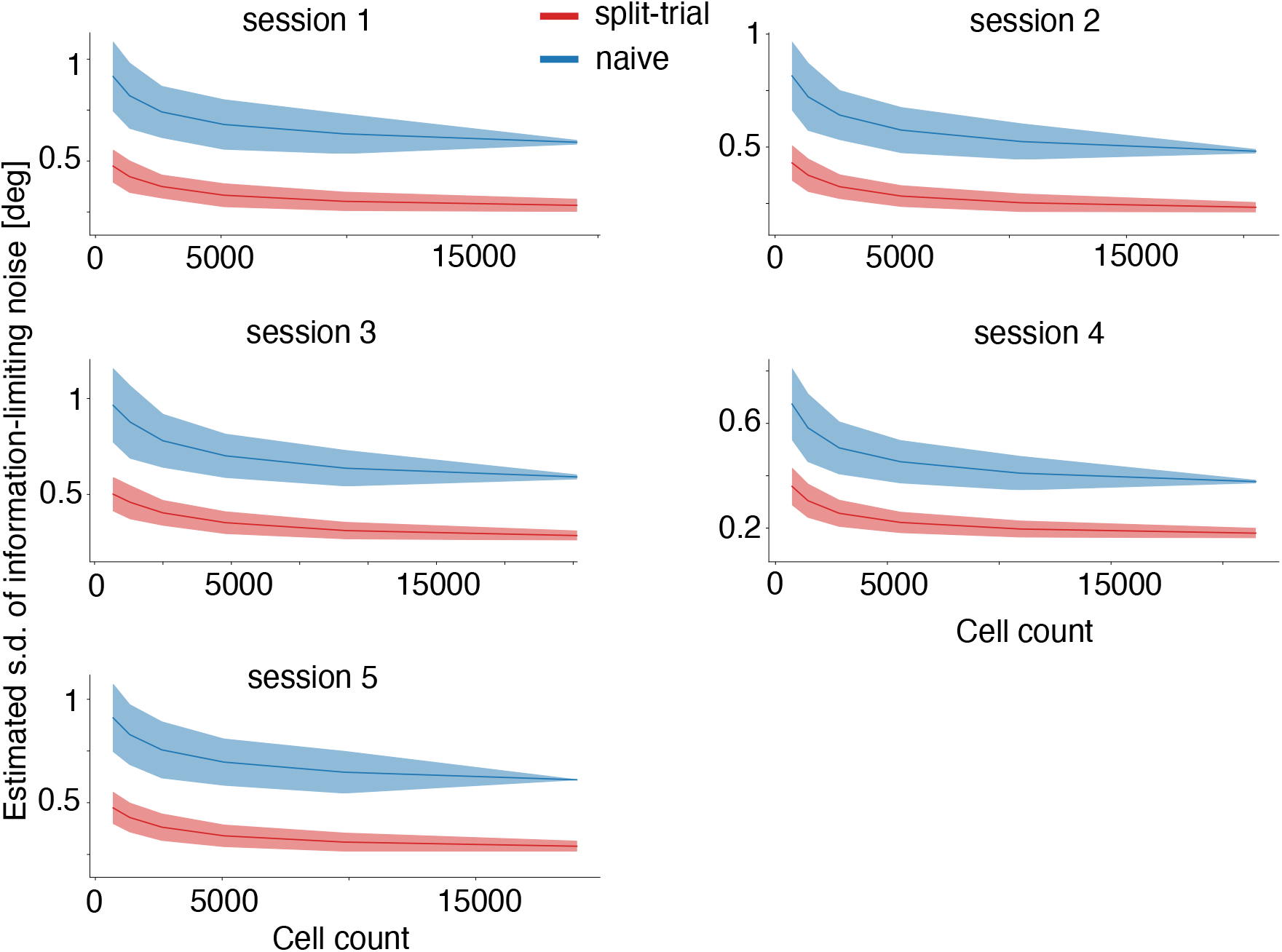
Results on the inferred standard deviation of information-limiting noise from the densely sampled orientation condition. Red: estimates from split-trial analysis. Blue: estimates from the direct (naive) decoding analysis.

**Figure S12:**
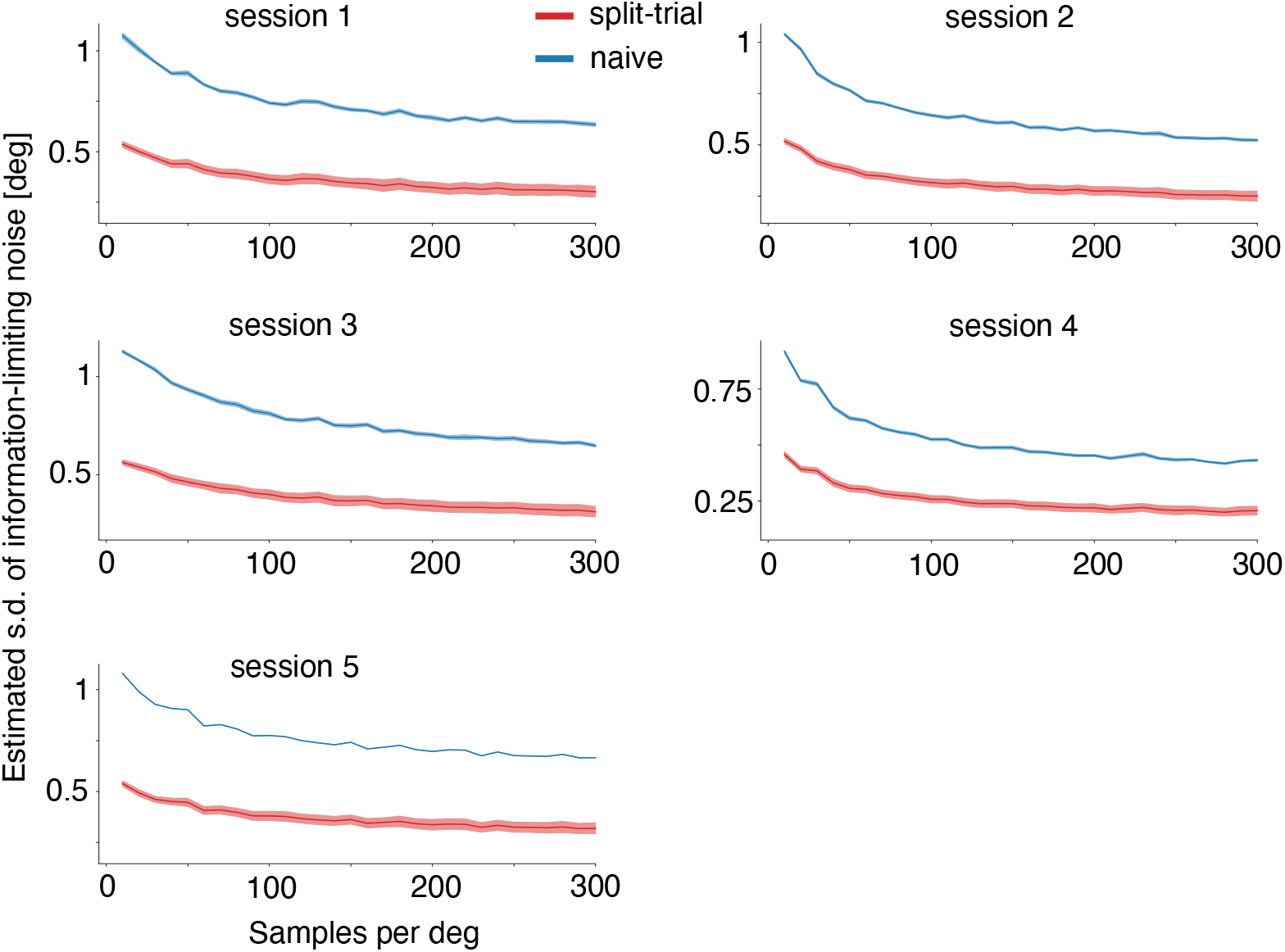
Results on the inferred standard deviation of information-limiting noise from the densely sampled orientation condition when the dataset was downsampled. Data from the dense discrimination sessions was downsampled to a range of different stimulus densities, up to 300, which is about the full dataset density. Notably, the split-trial estimates are stable and no longer appear to decrease after ∼150 samples/deg, suggesting that additional experimental trials would not affect the estimate.

**Figure S13:**
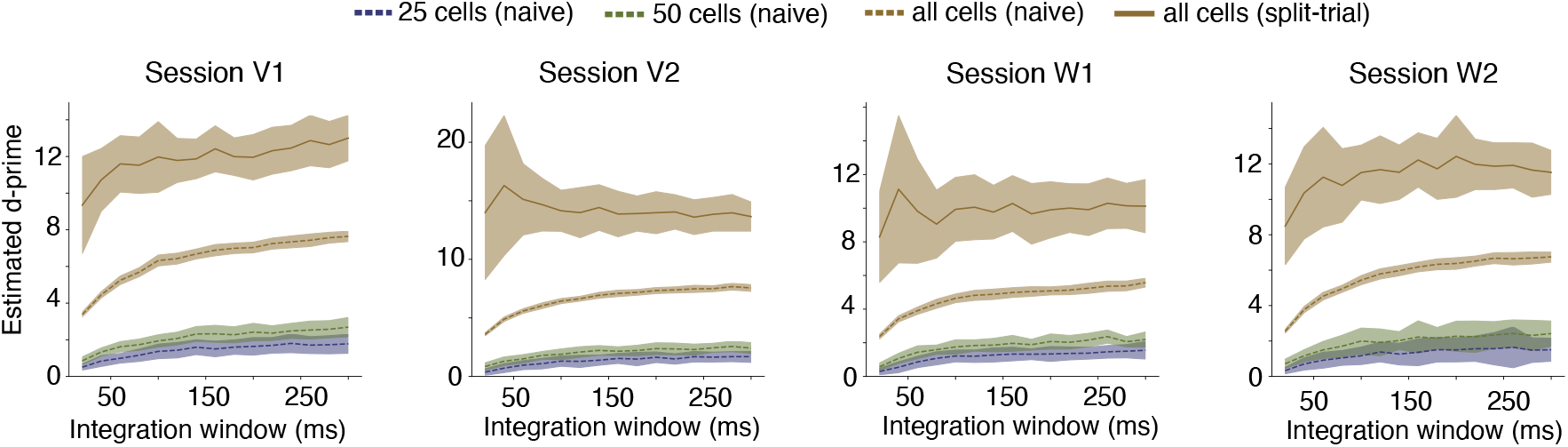
Results on how d-prime values scale with the increasing integration windows. Results from the split-trial analysis shown as solid line, showing relatively little change due to temporal integration. The four sessions have 770, 828, 509, 529 neurons, respectively.

**Figure S14:**
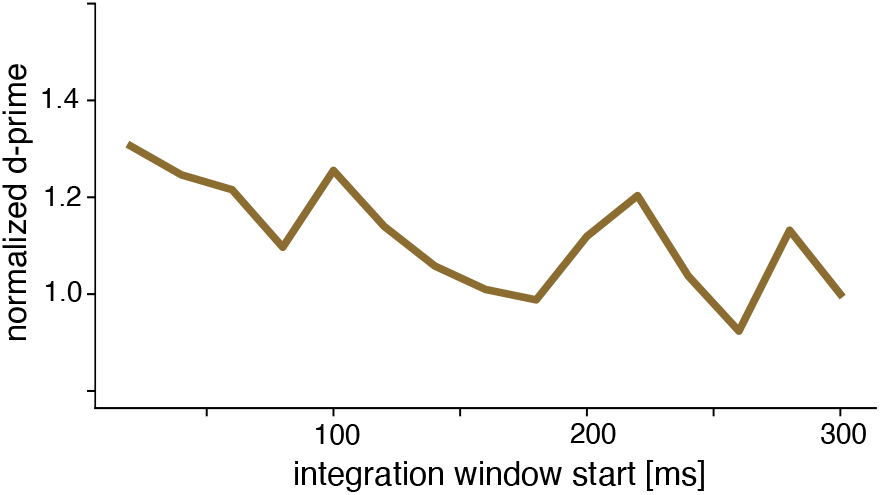
Average normalized d-prime values across normalized sessions (n=4) when windows are 20ms wide and discrete steps are taken through the 300ms window. D-prime values were inferred using split-trial analysis based on all neurons from individual sessions. The results show a trend that indicates information-limiting noise increases as reward delivery time approaches.

**Figure S15:**
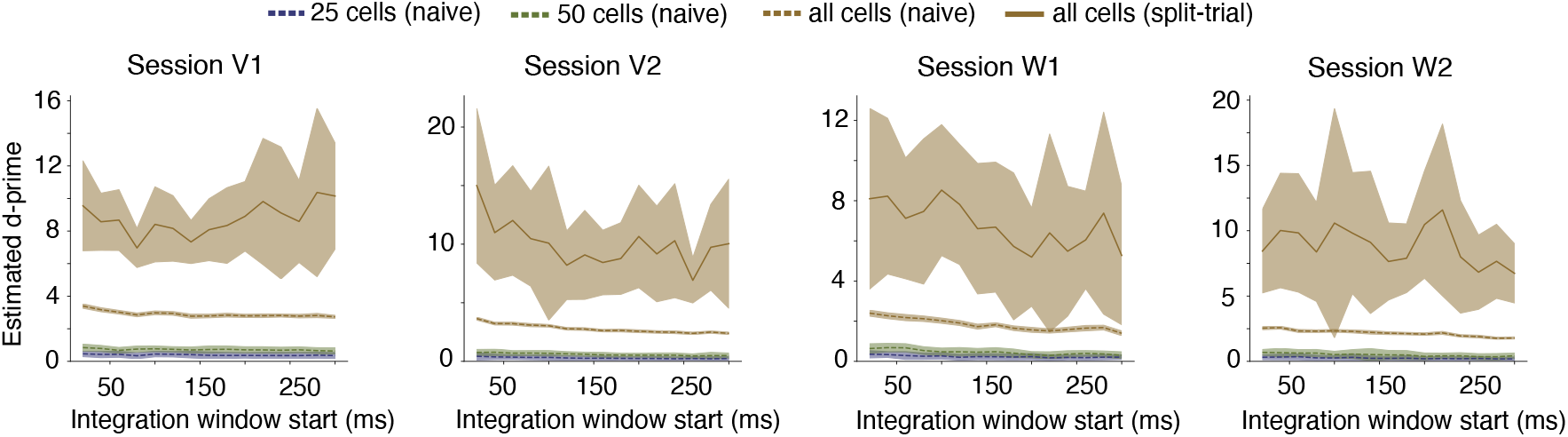
Non-normalized version of discrete windows individual sessions from the dataset from [25]. Despite early windows having higher values than later windows, they are typically lower than the (final) values recovered from the integrated calculations.

## References

[1] Ehud Zohary, Michael N Shadlen, and William T Newsome. Correlated neuronal discharge rate and its implications for psychophysical performance. Nature, 370(6485):140, 1994.

[2] Bruno B Averbeck, Peter E Latham, and Alexandre Pouget. Neural correlations, population coding and computation. Nature Reviews Neuroscience, 7(5):358–366, 2006.

[3] Marlene R Cohen and John HR Maunsell. Attention improves performance primarily by reducing interneuronal correlations. Nature neuroscience, 12(12):1594, 2009.

[4] Stefano Panzeri, Monica Moroni, Houman Safaai, and Christopher D Harvey. The structures and functions of correlations in neural population codes. Nature Reviews Neuroscience, 23(9):551–567, 2022.

[5] Rubén Moreno-Bote, Jeffrey Beck, Ingmar Kanitscheider, Xaq Pitkow, Peter Latham, and Alexandre Pouget. Information-limiting correlations. Nature Neuroscience, 17(10):1410–1417, 2014.

[6] Ingmar Kanitscheider, Ruben Coen-Cagli, and Alexandre Pouget. Origin of information-limiting noise correlations. Proceedings of the National Academy of Sciences, 112(50):E6973–E6982, 2015.

[7] Marlene R Cohen and Adam Kohn. Measuring and interpreting neuronal correlations. Nature neuroscience, 14(7):811, 2011.

[8] Jude F Mitchell, Kristy A Sundberg, and John H Reynolds. Spatial attention decorrelates intrinsic activity fluctuations in macaque area v4. Neuron, 63(6):879–888, 2009.

[9] Yong Gu, Sheng Liu, Christopher R Fetsch, Yun Yang, Sam Fok, Adhira Sunkara, Gregory C DeAngelis, and Dora E Angelaki. Perceptual learning reduces interneuronal correlations in macaque visual cortex. Neuron, 71(4):750–761, 2011.

[10] Martin Vinck, Renata Batista-Brito, Ulf Knoblich, and Jessica A Cardin. Arousal and locomotion make distinct contributions to cortical activity patterns and visual encoding. Neuron, 86(3):740–754, 2015.

[11] Akash Umakantha, Rudina Morina, Benjamin R Cowley, Adam C Snyder, Matthew A Smith, and M Yu Byron. Bridging neuronal correlations and dimensionality reduction. Neuron, 109(17):2740– 2754, 2021. PMCID: PMC8505167.

[12] Ingmar Kanitscheider, Ruben Coen-Cagli, Adam Kohn, and Alexandre Pouget. Measuring fisher information accurately in correlated neural populations. PLoS computational biology, 11(6):e1004218, 2015.

[13] Xaq Pitkow, Sheng Liu, Dora E Angelaki, Gregory C DeAngelis, and Alexandre Pouget. How can single sensory neurons predict behavior? Neuron, 87(2):411–423, 2015.

[14] Guillaume Hennequin, Yashar Ahmadian, Daniel B Rubin, Máté Lengyel and Kenneth D Miller. The dynamical regime of sensory cortex: stable dynamics around a single stimulus-tuned attractor account for patterns of noise variability. Neuron, 98(4):846–860, 2018.

[15] Larry F Abbott and Peter Dayan. The effect of correlated variability on the accuracy of a population code. Neural computation, 11(1):91–101, 1999.

[16] Alexander S Ecker, George H Denfield, Matthias Bethge, and Andreas S Tolias. On the structure of neuronal population activity under fluctuations in attentional state. Journal of Neuroscience, 36(5):1775–1789, 2016.

[17] Maoz Shamir and Haim Sompolinsky. Implications of neuronal diversity on population coding. Neural computation, 18(8):1951–1986, 2006.

[18] Felix Franke, Michele Fiscella, Maksim Sevelev, Botond Roska, Andreas Hierlemann, and Rava Azeredo da Silveira. Structures of neural correlation and how they favor coding. Neuron, 89(2):409– 422, 2016.

[19] Joel Zylberberg, Jon Cafaro, Maxwell H Turner, Eric Shea-Brown, and Fred Rieke. Directionselective circuits shape noise to ensure a precise population code. Neuron, 89(2):369–383, 2016.

[20] Alexander S Ecker, Philipp Berens, Andreas S Tolias, and Matthias Bethge. The effect of noise correlations in populations of diversely tuned neurons. The Journal of Neuroscience, 31(40):14272– 14283, 2011. PMCID: PMC3221941.

[21] Rava Azeredo da Silveira and Fred Rieke. The geometry of information coding in correlated neural populations. Annual Review of Neuroscience, 44:403–424, 2021.

[22] Oleg I Rumyantsev, Jérôme A Lecoq, Oscar Hernandez, Yanping Zhang, Joan Savall, Radosław Chrapkiewicz, Jane Li, Hongkui Zeng, Surya Ganguli, and Mark J Schnitzer. Fundamental bounds on the fidelity of sensory cortical coding. Nature, 580(7801):100–105, 2020.

[23] Mohammad Mehdi Kafashan, Anna W Jaffe, Selmaan N Chettih, Ramon Nogueira, Iñigo Arandia-Romero, Christopher D Harvey, Rubén Moreno-Bote, and Jan Drugowitsch. Scaling of sensory information in large neural populations shows signatures of information-limiting correlations. Nature communications, 12(1):473, 2021.

[24] Klaus Wimmer, Duane Q Nykamp, Christos Constantinidis, and Albert Compte. Bump attractor dynamics in prefrontal cortex explains behavioral precision in spatial working memory. Nature neuroscience, 17(3):431–439, 2014.

[25] Ramon Bartolo, Richard C Saunders, Andrew R Mitz, and Bruno B Averbeck. Information-limiting correlations in large neural populations. Journal of Neuroscience, 40(8):1668–1678, 2020.

[26] Omer Hazon, Victor H Minces, David P Tomàs, Surya Ganguli, Mark J Schnitzer, and Pablo E Jercog. Noise correlations in neural ensemble activity limit the accuracy of hippocampal spatial representations. Nature communications, 13(1):4276, 2022.

[27] Carsen Stringer, Michalis Michaelos, Dmitri Tsyboulski, Sarah E Lindo, and Marius Pachitariu. High-precision coding in visual cortex. Cell, 184(10):2767–2778, 2021.

[28] Amin Nejatbakhsh, Isabel Garon, and Alex Williams. Estimating noise correlations across continuous conditions with wishart processes. Advances in Neural Information Processing Systems, 36:54032–54045, 2023.

[29] C Radhakrishna Rao et al. Information and the accuracy attainable in the estimation of statistical parameters. Bull. Calcutta Math. Soc, 37(3):81–91, 1945.

[30] Alexander S Ecker, Philipp Berens, R James Cotton, Manivannan Subramaniyan, George H Denfield, Cathryn R Cadwell, Stelios M Smirnakis, Matthias Bethge, and Andreas S Tolias. State dependence of noise correlations in macaque primary visual cortex. Neuron, 82(1):235–248, 2014.

[31] Neil C Rabinowitz, Robbe L Goris, Marlene Cohen, and Eero P Simoncelli. Attention stabilizes the shared gain of v4 populations. Elife, 4:e08998, 2015.

[32] David Marvin Green, John A Swets, et al. Signal detection theory and psychophysics, volume 1. Wiley New York, 1966.

[33] Zaki Ajabi, Alexandra T Keinath, Xue-Xin Wei, and Mark P Brandon. Population dynamics of the thalamic head direction system during drift and reorientation. bioRxiv, 2021.

[34] Zaki Ajabi, Alexandra T Keinath, Xue-Xin Wei, and Mark P Brandon. Population dynamics of head-direction neurons during drift and reorientation. Nature, 615(7954):892–899, 2023.

[35] Jeffrey S Taube, Robert U Muller, and James B Ranck. Head-direction cells recorded from the postsubiculum in freely moving rats. i. description and quantitative analysis. Journal of Neuroscience, 10(2):420–435, 1990.

[36] Jonathan Green, Atsuko Adachi, Kunal K Shah, Jonathan D Hirokawa, Pablo S Magani, and Gaby Maimon. A neural circuit architecture for angular integration in drosophila. Nature, 546(7656):101– 106, 2017.

[37] Daniel Turner-Evans, Stephanie Wegener, Herve Rouault, Romain Franconville, Tanya Wolff, Johannes D Seelig, Shaul Druckmann, and Vivek Jayaraman. Angular velocity integration in a fly heading circuit. Elife, 6:e23496, 2017.

[38] Kechen Zhang. Representation of spatial orientation by the intrinsic dynamics of the head-direction cell ensemble: a theory. Journal of Neuroscience, 16(6):2112–2126, 1996.

[39] A David Redish, Adam N Elga, and David S Touretzky. A coupled attractor model of the rodent head direction system. Network: computation in neural systems, 7(4):671, 1996.

[40] Rani Ben-Yishai, R Lev Bar-Or, and Haim Sompolinsky. Theory of orientation tuning in visual cortex. Proceedings of the National Academy of Sciences, 92(9):3844–3848, 1995.

[41] Xue-Xin Wei, Ding Zhou, Andres Grosmark, Zaki Ajabi, Fraser Sparks, Pengcheng Zhou, Mark Brandon, Attila Losonczy, and Liam Paninski. A zero-inflated gamma model for post-deconvolved calcium imaging traces. Neurons, Behavior, Data analysis, and Theory, 3(2):1–21, 2020.

[42] David H Hubel and Torsten N Wiesel. Receptive fields, binocular interaction and functional architecture in the cat’s visual cortex. The Journal of physiology, 160(1):106, 1962.

[43] Nikolaus Kriegeskorte and Xue-Xin Wei. Neural tuning and representational geometry. Nature Reviews Neuroscience, 22(11):703–718, 2021.

[44] Annemarie Wolff, Nareg Berberian, Mehrshad Golesorkhi, Javier Gomez-Pilar, Federico Zilio, and Georg Northoff. Intrinsic neural timescales: temporal integration and segregation. Trends in cognitive sciences, 26(2):159–173, 2022.

[45] Yuzhi Chen, Wilson S Geisler, and Eyal Seidemann. Optimal temporal decoding of neural population responses in a reaction-time visual detection task. Journal of Neurophysiology, 99(3):1366–1379, 2008.

[46] Robbe LT Goris, Corey M Ziemba, J Anthony Movshon, and Eero P Simoncelli. Slow gain fluctuations limit benefits of temporal integration in visual cortex. Journal of vision, 18(8):8–8, 2018.

[47] Jamie D Roitman and Michael N Shadlen. Response of neurons in the lateral intraparietal area during a combined visual discrimination reaction time task. Journal of neuroscience, 22(21):9475– 9489, 2002.

[48] Natalie Steinemann, Gabriel M Stine, Eric Trautmann, Ariel Zylberberg, Daniel M Wolpert, and Michael N Shadlen. Direct observation of the neural computations underlying a single decision. Elife, 12:RP90859, 2024.

[49] Amy M Ni, Douglas A Ruff, Joshua J Alberts, Jen Symmonds, and Marlene R Cohen. Learning and attention reveal a general relationship between population activity and behavior. Science, 359(6374):463–465, 2018.

[50] Joao Barbosa, Heike Stein, Rebecca L Martinez, Adrià Galan-Gadea, Sihai Li, Josep Dalmau, Kirsten Adam, Josep Valls-Solé, Christos Constantinidis, and Albert Compte. Interplay between persistent activity and activity-silent dynamics in the prefrontal cortex underlies serial biases in working memory. Nature Neuroscience, 23(8):1016–1024, 2020.

[51] Kechen Zhang, Iris Ginzburg, Bruce L McNaughton, and Terrence J Sejnowski. Interpreting neuronal population activity by reconstruction: unified framework with application to hippocampal place cells. Journal of neurophysiology, 79(2):1017–1044, 1998.

[52] Jianqing Fan. On the optimal rates of convergence for nonparametric deconvolution problems. The Annals of Statistics, pages 1257–1272, 1991.

[53] Charles A Rohde. Generalized inverses of partitioned matrices. Journal of the Society for Industrial and Applied Mathematics, 13(4):1033–1035, 1965.

[54] Erich Leo Lehmann and George Casella. Theory of point estimation. Springer, 1998.

[55] Peggy Seriès, Peter E Latham, and Alexandre Pouget. Tuning curve sharpening for orientation selectivity: coding efficiency and the impact of correlations. Nature neuroscience, 7(10):1129, 2004.

[56] Steve Cherry. Singular value decomposition analysis and canonical correlation analysis. Journal of Climate, 9(9):2003–2009, 1996.

[57] Tony Cai, Jianqing Fan, and Tiefeng Jiang. Distributions of angles in random packing on spheres. The Journal of Machine Learning Research, 14(1):1837–1864, 2013.

